# A meta-analysis of epigenome-wide association studies in Alzheimer’s disease highlights novel differentially methylated loci across cortex

**DOI:** 10.1101/2020.02.28.957894

**Authors:** Rebecca G. Smith, Ehsan Pishva, Gemma Shireby, Adam R. Smith, Janou A.Y. Roubroeks, Eilis Hannon, Gregory Wheildon, Diego Mastroeni, Gilles Gasparoni, Matthias Riemenschneider, Armin Giese, Andrew J. Sharp, Leonard Schalkwyk, Vahram Haroutunian, Wolfgang Viechtbauer, Daniel L.A. van den Hove, Michael Weedon, Danielle Brokaw, Paul T. Francis, Alan J Thomas, Seth Love, Kevin Morgan, Jörn Walter, Paul D. Coleman, David A. Bennett, Philip L. De Jager, Jonathan Mill, Katie Lunnon

**Affiliations:** University of Exeter Medical School, College of Medicine and Health, University of Exeter, Exeter, UK; Department of Psychiatry and Neuropsychology, School for Mental Health and Neuroscience (MHeNS), Maastricht University, Maastricht, The Netherlands; Banner ASU Neurodegenerative Research Center, Biodesign Institute, Arizona State University, Tempe, Arizona, USA; Department of Genetics, University of Saarland (UdS), Saarbruecken, Germany; Department of Psychiatry and Psychotherapy, Saarland University Hospital (UKS), Homburg, Germany; Center for Neuropathology and Prion Research, Ludwig-Maximilians-University (LMU), Munich, Germany; Department of Genetics and Genomic Sciences, Icahn School of Medicine at Mount Sinai, New York, USA; School of Biological Sciences, University of Essex, Colchester, UK; Department of Psychiatry, The Icahn School of Medicine at Mount Sinai, New York, USA; Department of Neuroscience, The Icahn School of Medicine at Mount Sinai, New York, USA; JJ Peters VA Medical Center, Bronx, New York, USA; Laboratory of Translational Neuroscience, Department of Psychiatry, Psychosomatics and Psychotherapy, University of Wuerzburg, Würzburg, Germany; Institute of Neuroscience, Newcastle University, Newcastle Upon Tyne, UK; Dementia Research Group, Institute of Clinical Neurosciences, School of Clinical Sciences, University of Bristol, Bristol, UK; Human Genetics Group, University of Nottingham, Nottingham, UK; Rush Alzheimer’s Disease Center, Rush University Medical Center, Chicago, IL, USA; Center for Translational and Computational Neuroimmunology, Department of Neurology and Taub Institute, Columbia University Medical Center, New York, USA; The Broad Institute of MIT and Harvard, Cambridge, Massachusetts, USA

## Abstract

Epigenome-wide association studies of Alzheimer’s disease have highlighted neuropathology-associated DNA methylation differences, although existing studies have been limited in sample size and utilized different brain regions. Here, we combine data from six DNA methylomic studies of Alzheimer’s disease (N=1,453 unique individuals) to identify differential methylation associated with Braak stage in different brain regions and across cortex. We identify 236 CpGs in the prefrontal cortex, 95 CpGs in the temporal gyrus and ten CpGs in the entorhinal cortex at Bonferroni significance, with none in the cerebellum. Our cross-cortex meta-analysis (N=1,408 donors) identifies 220 CpGs associated with neuropathology, annotated to 121 genes, of which 84 genes have not been previously reported at this significance threshold. We have replicated our findings using two further DNA methylomic datasets consisting of a further > 600 unique donors. The meta-analysis summary statistics are available in our online data resource (www.epigenomicslab.com/ad-meta-analysis/).

## INTRODUCTION

Alzheimer’s disease (AD) is a chronic neurodegenerative disease that is accompanied by memory problems, confusion and changes in mood, behavior and personality. AD accounts for ~60% of dementia cases, which affected 43.8 million people worldwide in 2016^1^. The disease is defined by two key pathological hallmarks in the brain: extracellular plaques comprised of amyloid-beta protein and intracellular neurofibrillary tangles of hyperphosphorylated tau protein^2–4^. These neuropathological changes are thought to occur perhaps decades before clinical symptoms manifest and the disease is diagnosed^4^. AD is a multi-factorial and complex disease, with the risk of developing disease still largely unknown despite numerous genetic and epidemiological studies over recent years.

Several studies have suggested that epigenetic mechanisms may play a role in disease etiology. In recent years a number of epigenome-wide association studies (EWAS) have been performed in AD brain samples, which have predominantly utilized the Illumina Infinium HumanMethylation450K BeadChip (450K array) in conjunction with bisulfite-treated DNA to assess levels of DNA methylation in cortical brain tissue from donors with varying degrees of AD pathology^5–12^. Independently these studies have identified a number of loci that show robust differential DNA methylation in disease, and many of these overlap between studies, for example loci annotated to *ANK1, RHBDF2, HOXA3, CDH23* and *RPL13* have been consistently reported.

Here we have performed a meta-analysis of six independent existing EWAS of AD brain^5–8,10,12^, totalling 1,453 independent donors, to identify robust and consistent differentially methylated loci associated with Braak stage, used as a measure of neurofibrillary tangle spread through the brain, in different brain regions and across the cortex. In our intra-tissue meta-analysis we identify 236 CpGs in the prefrontal cortex (N = 959 samples), 95 in the temporal gyrus (N = 608 samples) and ten in the entorhinal cortex (N = 189 samples) at Bonferroni significance, with none in the cerebellum (N = 533 samples). Our cross-cortex meta-analysis (N = 1,408 individuals) identified 220 Bonferroni significant CpGs, which were replicated in two further independent DNA methylation datasets. Our meta-analysis approach provides additional power to detect DNA methylomic variation associated with AD pathology at novel loci, in addition to providing further replication of loci that have been previously identified in the smaller independent EWAS.

## RESULTS

### Pathology-associated DNA methylation signatures in discrete cortical brain regions

We identified six EWAS of DNA methylation in AD that had been generated using the 450K array and had a cohort size of > 50 unique donors. All had data on Braak stage available, which we used as a standardized measure of tau pathology spread through the brain (Table 1). We were interested in identifying epigenomic profiles associated with Braak stage in specific brain regions, leveraging additional power by meta-analysing multiple studies to identify novel loci. To this end, we performed an EWAS in each available tissue and cohort separately, looking for an association between DNA methylation and Braak stage, whilst controlling for age and sex (all tissues) and neuron/glia proportion (cortical bulk tissues only), with surrogate variables added as appropriate to reduce inflation. For discovery, we then used the estimated effect size (ES) and standard errors (SEs) from these six studies (N = 1,453 unique donors) for a fixed-effect inverse variance weighted meta-analysis separately for each tissue (prefrontal cortex: three cohorts, N = 959; temporal gyrus: four cohorts, N = 608, entorhinal cortex: two cohorts, N = 189 cerebellum: four cohorts, N = 533) (Supplementary Figure 1).

**Table 1:** Demographic information for cohorts included in the meta-analyses. Sample numbers, split of males (M)/females (F) and mean age at death in years (± standard deviation [SD]) are shown for individuals with low pathology (Braak 0-II), mid-stage pathology (Braak III-IV) and severe pathology (Braak V-VI) in each cohort. Shown are the bulk tissues available from each cohort, which included the cerebellum, entorhinal cortex, middle temporal gyrus, prefrontal cortex and superior temporal gyrus. In the discovery meta-analyses, we used data from six EWAS using the 450K array, which all had > 50 unique donors. For replication we used two cohorts. The Munich cohort had 450K data from bulk prefrontal cortex tissue, as well as data available from sorted neuronal and non-neuronal cell populations from the occipital cortex. The BDR cohort had EPIC array data available from bulk prefrontal cortex samples. For the meta-analyses, superior temporal gyrus and middle temporal gyrus samples were both classed as temporal gyrus samples. Shown are final numbers for all cohorts after data quality control. Ancestry is reported for the discovery cohorts and is the number of unique individuals that had the following inferred ethnicities from the 1000 genomes reference panel: European (Eu), African (Af), American (Am), East Asian (As).

| Stage | Cohort | Unique individuals | Ancestry (Eu/Af/Am/As) | Braak | Number | Sex (M/F) | Age at death in (± SD) | Tissues analysed |
| --- | --- | --- | --- | --- | --- | --- | --- | --- |
| DISCOVERY | London 1 | 113 | 112/0/1/0 | 0-II | 29 | 13/16 | 77.6 (12.8) | Prefrontal cortex, entorhinal cortex, superior temporal gyrus, cerebellum (Bulk) |
|  |  |  |  | III-IV | 18 | 7/11 | 88.5 (5.2) |  |
|  |  |  |  | V-VI | 66 | 26/40 | 85.4 (8.1) |  |
|  | London 2 | 95 | 92/1/2/0 | 0-II | 23 | 12/11 | 76.1 (10.0) | Entorhinal cortex, cerebellum (Bulk) |
|  |  |  |  | III-IV | 16 | 3/13 | 87.6 (6.4) |  |
|  |  |  |  | V-VI | 56 | 26/30 | 81.5 (8.6) |  |
|  | Mount Sinai | 146 | 113/20/11/2 | 0-II | 60 | 32/28 | 82 (7.6) | Prefrontal cortex, superior temporal gyrus (Bulk) |
|  |  |  |  | III-IV | 42 | 12/30 | 88.8 (6.6) |  |
|  |  |  |  | V-VI | 44 | 12/32 | 88.0 (7.5) |  |
|  | Arizona 1 | 302 | 302/0/0/0 | 0-II | 61 | 40/21 | 80.3 (8.2) | Middle temporal gyrus, cerebellum (Bulk) |
|  |  |  |  | III-IV | 97 | 50/47 | 86.9 (6.9) |  |
|  |  |  |  | V-VI | 144 | 63/81 | 82.3 (8.5) |  |
|  | Arizona 2 | 88 | 88/0/0/0 | 0-II | 16 | 10/6 | 82.5 ( 5.0) | Middle temporal gyrus, cerebellum (Bulk) |
|  |  |  |  | III-IV | 45 | 21/24 | 86.7 (5.1) |  |
|  |  |  |  | V-VI | 27 | 12/15 | 84.6 (7.1) |  |
|  | ROS/MAP | 709 | 709/0/0/0 | 0-II | 143 | 70/73 | 83.2 (6.0) | Prefrontal cortex (Bulk) |
|  |  |  |  | III-IV | 409 | 144/266 | 86.9 (4.1) |  |
|  |  |  |  | V-VI | 157 | 45/113 | 87.8 (3.5) |  |
| REPLICATION | Munich | 45 | - | 0-II | 9 | 5/4 | 76.7 (10.9) | Prefrontal cortex (Bulk) |
|  |  |  |  | III-IV | 7 | 1/6 | 82.1 (5.2) |  |
|  |  |  |  | V-VI | 29 | 12/17 | 79.2 (8.5) |  |
|  |  | 26 | - | 0-II | 11 | 7/4 | 75.9 (8.5) | Occipital cortex (Sorted cells) |
|  |  |  |  | III-IV | 5 | 1/4 | 85.0 (6.5) |  |
|  |  |  |  | V-VI | 10 | 4/6 | 77.9 (6.6) |  |
|  | BDR | 590 | - | 0-II | 196 | 100/96 | 83.6 (10.6) | Prefrontal cortex (Bulk) |
|  |  |  |  | III-IV | 136 | 91/65 | 85.1 (7.45) |  |
|  |  |  |  | V-VI | 258 | 128/130 | 82.5 (8.5) |  |

The prefrontal cortex represented our largest dataset (N = 959 samples) and we identified 236 Bonferroni significant differentially methylated positions (DMPs) (P < 1.238 x 10^-7^ to account for 403,763 probes), of which 193 were annotated to 137 genes, with 43 unannotated loci based on Illumina UCSC annotation (Figure 1a, Supplementary Figure 2, Supplementary Data 1). Previous EWAS of the prefrontal cortex have consistently reported the *HOXA* gene cluster as a region that is hypermethylated in AD^6,7^, with a cell-type specific EWAS demonstrating this is neuronal-derived^11^. Indeed, the most significant DMP in the prefrontal cortex in our meta-analysis resided in *HOXA3* (cg22962123: ES [defined as the methylation difference between Braak 0 and Braak VI] = 0.042, P = 5.97 x 10^-15^), with a further 16 of the Bonferroni significant DMPs also annotated to this gene. This locus appeared to be particularly hypermethylated with higher Braak stage in the prefrontal cortex, and to a slightly lesser extent in the temporal gyrus (Supplementary Figure 3). There was no significant difference in methylation at this locus in the entorhinal cortex (P = 0.864), which is interesting given that the entorhinal cortex may succumb to pathology early in the disease process (Braak stage III). Of the 236 prefrontal cortex DMPs, 92% (217 probes) were nominally significant (P < 0.05) in the temporal gyrus, of which 12% (28 probes) were Bonferroni significant, whilst 9% (22 probes) were nominally significant in the entorhinal cortex, with 1% (3 probes) reaching Bonferroni significance (Figure 1b). The effect sizes for the 236 Bonferroni significant prefrontal cortex DMPs were correlated with the effect sizes for the same probes in both the temporal gyrus (Pearson’s correlation coefficient (r) = 0.94, P = 6.17 x 10^-112^) and entorhinal cortex (r = 0.58, P = 1.80 x 10^-22^) and were enriched for probes with the same direction of effect (sign test: temporal gyrus P = 5.07 x 10^-67^, entorhinal cortex P = 6.88 x 10^-26^) (Supplementary Figure 4). For the 236 Bonferroni significant prefrontal cortex DMPs these had the largest effect sizes in the prefrontal cortex, with a smaller effect size in the temporal gyrus and entorhinal cortex (Figure 1c). Of these 236 DMPs, 29 of these had being previously reported at Bonferroni significance in previous publications on the individual cohorts^5–7,12^, including one probe annotated to *ANK1*, one probe annotated to *HOXA3*, one probe annotated to *PPT2/PRRT1* and two probes annotated to *RHBDF2*, amongst others. However, our approach has identified 207 novel Bonferroni significant DMPs (although several had been reported in previous studies at a more relaxed significance threshold, or in regional analyses). This included several additional probes residing in genes already identified (from another probe) in earlier studies, for example a further 16 probes in *HOXA3* and two probes in *PPT2/PRRT1*. Interestingly, we also identified a number of novel genes, including some which featured multiple Bonferroni significant DMPs including for example seven probes in *AGAP2* and five probes in *SLC44A2*, amongst others. One other noteworthy novel Bonferroni significant DMP in the prefrontal cortex was cg08898775 (ES = 0.019, P = 4.03 x 10^-9^), annotated to *ADAM10*, which encodes for α-secretase which cleaves APP in the non-amyloidogenic pathway. A differentially methylated region (DMR) analysis, which allowed us to identify areas of the genome consisting of ≥ 2 DMPs, revealed 262 significant DMRs in the prefrontal cortex (Supplementary Data 2), the most significant containing 20 probes and located in *HOXA3* (chr7:27,153,212-27,155,234: Sidak-corrected p = 8.21 x 10^-50^, Supplementary Figure 5), as well as several other DMRs in the *HOXA* gene cluster.

**Figure 1:**
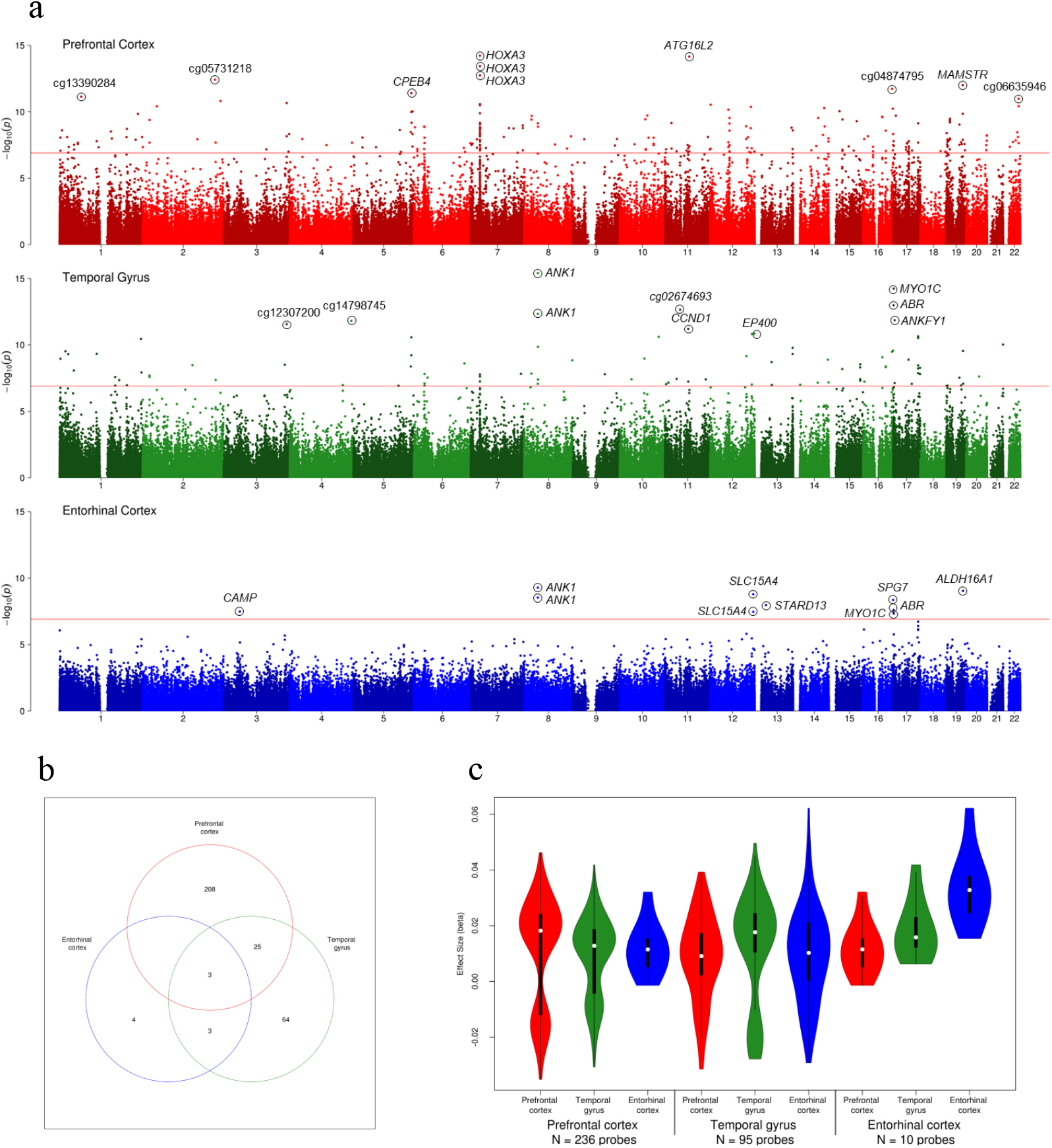
Intra-tissue meta-analyses of AD methylomic studies highlights Bonferroni significant differentially methylated positions (DMPs) in all cortical tissues. (**a**) A Manhattan plot for the prefrontal cortex (red, N = 959), temporal gyrus (green, N = 608) and entorhinal cortex (blue, N = 189) meta-analyses, with the ten most significant DMPs circled on the plot and Illumina UCSC gene name shown if annotated, or CpG ID if unannotated. The X-axis shows chromosomes 1-22 and the Y-axis shows −log10(p), with the horizontal red line denoting Bonferroni significance (P < 1.238 x 10^-7^). (**b**) A Venn diagram highlighting overlapping DMPs at Bonferroni significance across the cortical tissues. (**c**) In each cortical brain region the Bonferroni significant DMPs identified in that region usually had a greater effect size (ES) there, than in any of the other cortical regions. The X-axis represents the methylation (beta) ES between individuals that are Braak stage 0 and VI. Data is separated on the Y-axis by tissue analysis (large text) with the corresponding data at these probes in other tissues (small text). The white dot in the centre represents the median, the dark box represents the interquartile range (IQR), whilst the whisker lines represent the “minimum” (quartile 1 - 1.5 x IQR) and the “maximum” (quartile 3 + 1.5 x IQR). The coloured violin represents all samples including outliers, meaning that the “minimum” and “maximum” may not extend to the end of the violin.

A meta-analysis of temporal gyrus EWAS datasets (N = 608 samples) identified 95 Bonferroni significant probes, of which 75 were annotated to 53 genes, with 20 unannotated probes using Illumina UCSC annotation (Figure 1a, Supplementary Figure 6, Supplementary Data 3). The most significant probe was cg11823178 (ES = 0.029, P = 3.97 x 10^-16^, Supplementary Figure 7), which is annotated to the *ANK1* gene, with the fifth (cg05066959: ES = 0.042, P = 4.58 x 10^-13^) and 82^nd^ (cg16140558: ES = 0.013, P = 8.44 x 10^-8^) most significant probes in the temporal gyrus also being annotated to nearby CpGs in that gene. This locus has been widely reported to be hypermethylated in AD from prior EWAS^5,6,8,12^, as well as in other neurodegenerative diseases such as Huntington’s disease and Parkinson’s disease^13^. Another noteworthy gene is *RHBDF2*, where five Bonferroni significant DMPs in the temporal gyrus were annotated to (cg05810363: ES = 0.029, P = 2.25 x 10^-11^; cg13076843: ES = 0.031, P = 2.97 x 10^-11^; cg09123026: ES = 0.012, P = 3.46 x 10^-9^; cg12163800: ES = 0.025, P = 5.85 x 10^-9^; cg12309456: ES = 0.016, P = 1.33 x 10^-8^); and which has been highlighted in previous EWAS in AD in the individual cohorts^5,6,12^. Of the 95 Bonferroni significant DMPs in the temporal gyrus, 88% (84 probes) were nominally significant in the prefrontal cortex, of which 29% (28 probes) were Bonferroni significant, whilst 54% (51 probes) were nominally significant in the entorhinal cortex, of which 6% (6 probes) were Bonferroni significant (Figure 1b). Given the high degree of overlapping significant loci between the temporal gyrus and other cortical regions, it was not surprising that the ES of the 95 Bonferroni significant temporal gyrus probes were highly correlated with the ES of the same loci in both the prefrontal cortex (r = 0.91, P = 5.09 x 10^-38^) and entorhinal cortex (r = 0.77, P = 4.02 x 10^-20^) and were enriched for the same direction of effect (sign test: prefrontal cortex P = 5.05 x 10^-29^, entorhinal cortex = 2.30 x 10^-25^) (Supplementary Figure 8). The majority of the 95 Bonferroni significant DMPs in the temporal gyrus were hypermethylated, and the mean ES was greater in the temporal gyrus than the prefrontal cortex or entorhinal cortex (Figure 1c). Thirty-two of the 95 Bonferroni significant DMPs in the temporal gyrus have been previously reported to be significantly differentially methylated in published EWAS, including for example three probes in *ANK1* and the five probes in *RHBDF2*. Our meta-analysis approach in the temporal gyrus has identified 63 novel DMPs (at Bonferroni significance), including some novel genes with multiple DMPs, for example four probes in *RGMA* and two probes in *CCND1*, amongst others. Finally, our regional analysis highlighted 104 DMRs (Supplementary Data 4); the top DMR resided in the *ANK1* gene (chr8:41,519,308-41,519,399) and contained two probes (Sidak-corrected P = 1.72 x 10^-21^) (Supplementary Figure 9). The five DMPs in *RHBDF2* that we already highlighted also represented a significant DMR (Sidak-corrected P = 8.47 x 10^-21^), with three other genomic regions containing large, significant DMRs consisting of ≥ 10 probes, such as*MCF2L* (chr13:113698408-113699016 [10 probes], Sidak-corrected P = 1.16 x 10^-19^), *PRRT1/PPT2* (chr6:32120773-32121261 [17 probes], Sidak-corrected P = 4.90 x 10^-15^) and *HOXA5* (chr7:27184264-27184521 [10 probes], Sidak-corrected P = 1.60 x 10^-7^).

The final cortical region we had available was the entorhinal cortex (N = 189), where we identified ten Bonferroni significant probes in our meta-analysis, all of which were hypermethylated with higher Braak stage (Figure 1a, Supplementary Figure 10, Supplementary Data 5). These ten probes were annotated to eight genes (Illumina UCSC annotation), with two Bonferroni significant probes residing in each of the *ANK1* and *SLC15A4* genes. As with the temporal gyrus, the most significant DMP was cg11823178 (ES = 0.045, P = 5.22 x 10^-10^, Supplementary Figure 7), located within the *ANK1* gene, with the fourth most significant DMP being located within 100bp of that CpG (cg05066959: ES = 0.062, P = 2.93 x 10^-9^). In total, eight of the ten DMPs in the entorhinal cortex had been reported previously at Bonferroni significance, including the two probes in *ANK1*. Two of the Bonferroni significant DMPs we identified in the entorhinal cortex were novel CpGs (cg11563844: *STARD13*, cg04523589: *CAMP*), having not been reported as Bonferroni significant in previous EWAS. Of the ten entorhinal cortex probes, 90% (9 probes) were nominally significant in the temporal gyrus, of which 60% (6 probes) were Bonferroni significant, whilst 70% (7 probes) were nominally significant in the prefrontal cortex, of which 30% (3 probes) were Bonferroni significant (Figure 1b). Of the four DMPs that were Bonferroni significant in only the entorhinal cortex, three of these were nominally significant in at least one other tissue, with just one probe unique to the entorhinal cortex, annotated to *STARD13* (cg11563844, ES = 0.027, P = 1.07 x 10^-8^). The effect sizes of the ten Bonferroni significant DMPs in the entorhinal cortex were significantly correlated with the effect size of the same probes in the prefrontal cortex (r = 0.74, P = 0.01) and temporal gyrus (r = 0.85, P = 1.52 x 10^-3^) and were enriched for the same direction of effect (sign test: prefrontal cortex P = 0.021, temporal gyrus P = 1.95 x 10^-3^) (Supplementary Figure 11). The ten DMPs were hypermethylated in all three cortical regions, with the greatest Braak-associated ES in the entorhinal cortex (Figure 1c). A regional analysis identified seven DMRs (Supplementary Data 6); the top three DMRs *(RHBDF2:* chr17:74,475,240-74,475,402 [five probes], P = 7.68 x 10^-14^, Supplementary Figure 12; *ANK1:* chr8:41519308-41519399 [two probes], P = 4.89 x 10^-13^; *SLC15A4:* chr12:129281444-129281546 [three probes], P = 5.24 x 10^-12^) were significant in at least one of the other cortical regions we meta-analyzed.

To date, a few independent EWAS in AD have been undertaken in the cerebellum and none of these have reported any Bonferroni significant DMPs. In our meta-analysis we identified no Bonferroni significant DMPs, nor any DMRs in the cerebellum (Supplementary Figure 13), despite this analysis including 533 independent samples. There was no correlation of the ES for the Bonferroni significant DMPs we had identified in the meta-analyses of the three cortical regions with the ES of the same probes in the cerebellum (prefrontal cortex: r = 0.11, P = 0.08; temporal gyrus: r = 0.14, P = 0.17; entorhinal cortex: r = 0.48, P = 0.16; Supplementary Figure 14).

### 220 CpGs are differentially methylated across the cortex in AD

We were interested in combining data from across the different cortical tissues to identify common differentially methylated loci across the cortex and also to provide more power by utilizing data from 1,408 unique individuals with cortical EWAS data available. As multiple cortical tissues were available for some cohorts, a mixed-effects model was utilized. In this analysis we controlled for age, sex and neuron/glia proportion, with surrogate variables added as appropriate to reduce inflation. Using this approach, we identified 220 Bonferroni significant probes, of which 168 were annotated to 121 genes, with 52 DMPs unannotated using Illumina UCSC annotation (Figure 2a, Figure 2b, Table 2, Supplementary Data 7, Supplementary Figure 15). All of the 220 probes were nominally significant (P < 0.05) in ≥ two cohorts, with ten of these probes being nominally significant in all six cohorts (Supplementary Figure 16), which included single probes annotated to *ANK1, ABR, SPG7* and *WDR81*, two probes in *DUSP27*, three probes in *RHBDF2* and one unannotated probe. We observed similar DNA methylation patterns across all cortical cohorts and tissues for the 220 probes with 219 of the 220 DMPs showing the same direction of effect in at least five cohorts. In total, 154 of the DMPs were hypermethylated, with 66 hypomethylated, representing an enrichment for hypermethylation (P = 4.85 x 10^-10^). This pattern of methylation was evident across all cortical tissues but was not seen in the cerebellum (Supplementary Figure 17). Of the 220 DMPs we identified, 46 of these have been previously reported at Bonferroni significance in published EWAS, including multiple previously identified probes in *ANK1* (cg05066959, cg11823178), *MCF2L* (cg07883124, cg09448088), *PCNT* (cg00621289, cg04147621, cg23449541) and *RHBDF2* (cg05810363, cg12163800, cg12309456, cg13076843). The most significant probe we identified in our cross-cortex analysis was cg12307200 (Table 2, ES = −0.015, P = 4.48 x 10^-16^), which is intergenic and found at chr3:188664632, located between the *TPRG1* and *LPP* genes and had been previously reported at Bonferroni significance by De Jager and colleagues with respect to neuritic plaque burden^6^ and by Brokaw and colleagues with respect to post-mortem diagnosis^12^. Our cross-cortex meta-analysis approach has identified 174 novel DMPs (at Bonferroni significance), annotated to 102 genes. Although 11 of these genes had previously been reported at Bonferroni significance (another probe within that gene), the remaining 96 genes represent robust novel loci in AD. Many of these novel differentially methylated genes had multiple Bonferroni significant probes, for example five probes in *AGAP2*, three probes in *HOXB3* and *SLC44A2*, and two probes in *CDH9, CPEB4, DUSP27, GCNT2, MAMSTR, PTK6, RGMA, RHOB, SMURF1, THBS1, ZNF238* and *ZNF385A* (Supplementary Data 7). Although some of these loci may have been reported in earlier AD EWAS, none of these were at Bonferroni significance and so here represent robust novel loci.

**Figure 2:**
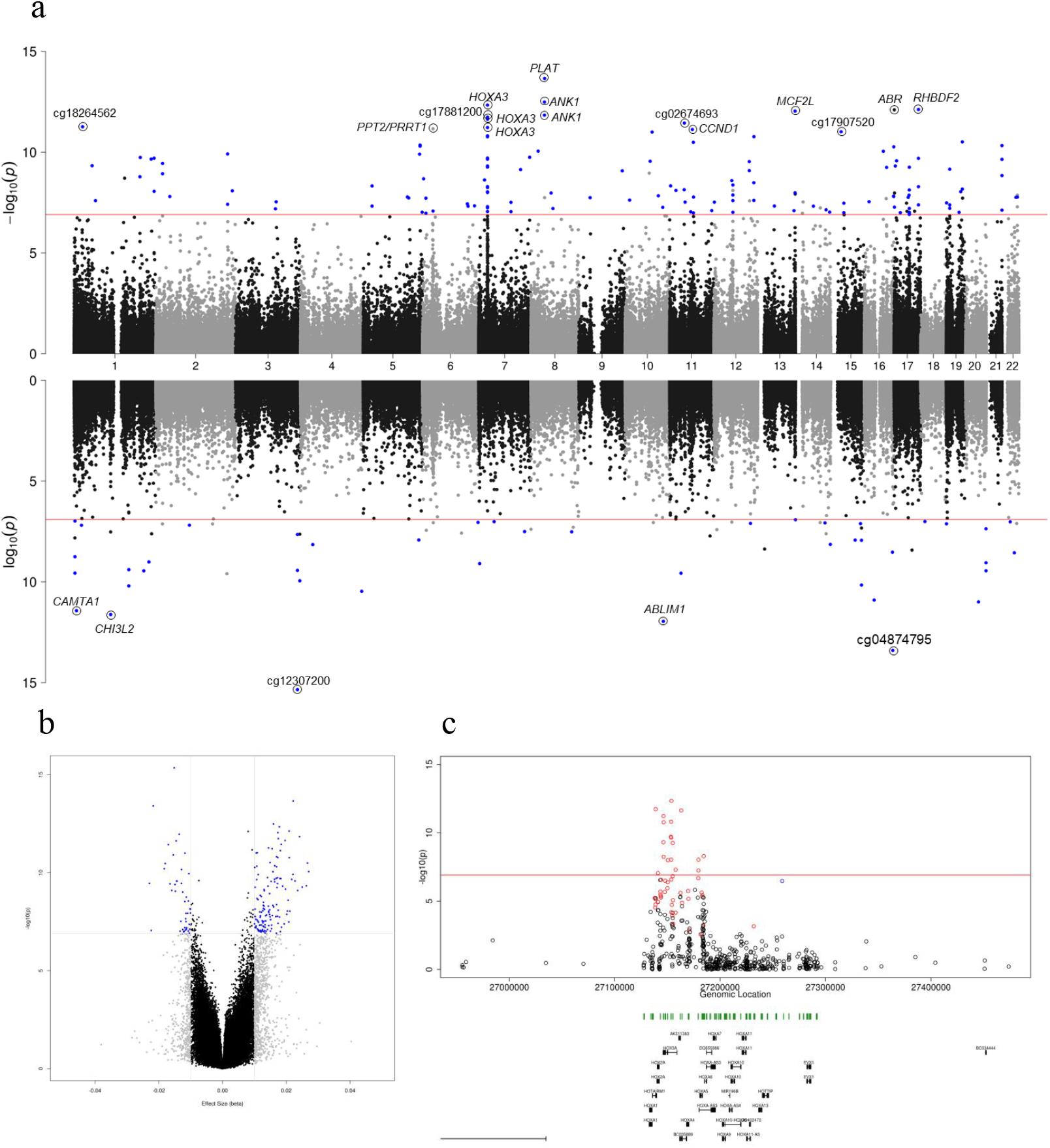
A cross-cortex meta-analysis identifies 220 Bonferroni significant differentially methylated positions (DMPs) associated with Braak stage. (**a**) A Miami plot of the cross-cortex meta-analyses (N = 1,408). Probes shown above the X-axis indicate hypermethylation with higher Braak stage, whilst probes shown below the X-axis indicate hypomethylation with higher Braak stage. The chromosome and genomic position are shown on the X-axis. The Y-axis shows −log10(p). The red horizontal lines indicate the Bonferroni significance level of P < 1.238 x 10^-7^. Probes with a methylation (beta) effect size (ES: difference between Braak 0-Braak VI) ≥ 0.01 and P < 1.238 x 10^-7^ are shown in blue. The 20 most significant DMPs are circled on the plot and Illumina UCSC gene name is shown if annotated, or CpG ID if unannotated. Exact p-values can be found in Table 2 and Supplementary Data 7. (**b**) A volcano plot showing the ES (X-axis) and −log10(p) (Y-axis) for the cross-cortical meta-analysis results. Gray probes indicate an ES between ≥ 0.01, whilst blue probes indicate an ES ≥ 0.01 and P < 1.238 x 10^-7^. (**c**) The most significant cross-cortex differentially methylated region (DMR) (chr7:27153212-27154305) contained 11 probes and resided in the *HOXA* region. The horizontal red line denotes the Bonferroni significance level of P < 1.238 x 10^-7^. Red probes represent a positive ES ≥ 0.01, blue probes represent a negative ES ≥ 0.01. Underneath the gene tracks are shown in black with CpG islands in green.

**Table 2:**
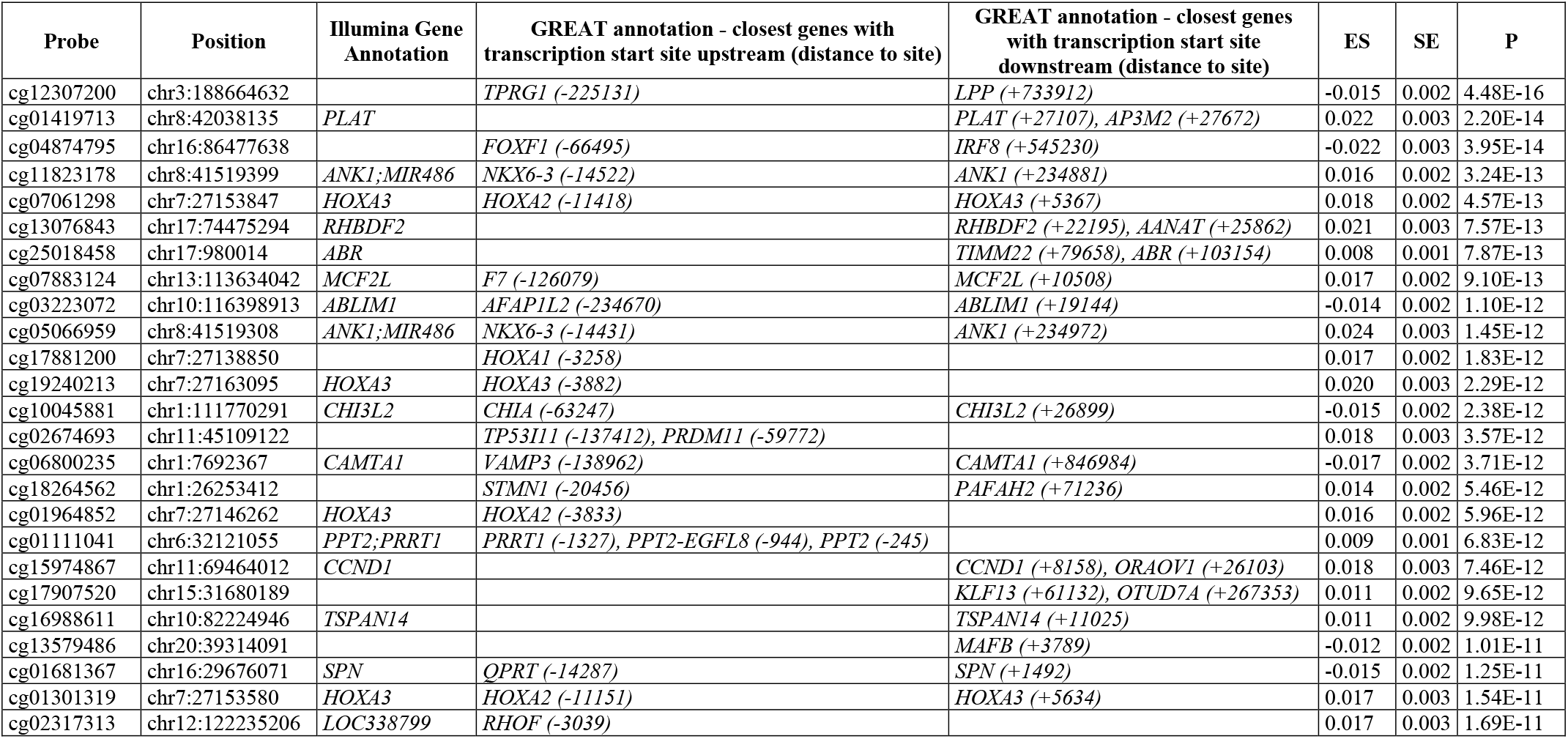
The 25 most significant differentially methylated positions (DMPs) associated with Braak stage from the cross-cortex inverse variance fixed effects meta-analysis. Probe information is provided corresponding to chromosomal location (hg19/GRCh37 genomic annotation), Illumina gene annotation, closest genes with a transcription start site upstream or downstream (from GREAT annotation). Shown for each DMP is the methylation (beta) effect size (ES), standard error (SE) and corresponding unadjusted P value from the inverse variance fixed effects meta-analysis model in the cross-cortex data (N = 1,408). All ES and SE have been multiplied by six to demonstrate the difference between Braak stage 0 and Braak stage VI samples. A more comprehensive table is provided in Supplementary Data 7.

We were interested to investigate whether specific functional pathways were differentially methylated in AD cortex and so performed a gene ontology pathway analysis of the 121 genes annotated to the 220 Bonferroni significant cross-cortex DMPs. We highlighted epigenetic dysfunction in numerous pathways (at nominal significance), interestingly including a number of developmental pathways, mainly featuring the *HOXA* and *HOXB* gene clusters (Supplementary Data 8). Given that we identified multiple DMPs in some genes, we were interested to investigate the correlation structure between probes in close proximity to each other to establish how many independent signals we had identified. Using a method developed to identify single nucleotide polymorphisms (SNPs) in linkage disequilibrium (LD)^14^, we collapsed the 220 Bonferroni significant loci into 165 independent (non-highly correlated [r < 0.6 over 1mb]) signals (Supplementary Data 9). We found that the largest reduction in signals occurred in the *HOXA* and *HOXB* gene clusters, with the 18 DMPs in the *HOXA* region representing only two independent signals, whilst the seven DMPs in the *HOXB* region represented one independent signal. Next we undertook a formal regional analysis to identify genomic regions of multiple adjacent DMPs and identified 221 DMRs, with the top DMR containing 11 probes and covering the *HOXA* region (chr7:27,153,212-27,154,305: P = 3.84 x 10^-35^) (Figure 2c, Supplementary Data 10). The *HOXA* gene cluster further featured a number of times in our DMR analysis; four of the ten most significant DMRs fell in this genomic region, including DMRs spanning four probes (chr7:27146237-27146445: P = 4.11 x 10^-27^), 33 probes (chr7:27183133-27184667: P = 2.22 x 10^-20^) and ten probes (chr7:27143235-27143806: P = 1.75 x 10^-18^).

### Replication of pathology associated DMPs in the cortex

To replicate our findings and to determine the cellular origin of DNA methylomic differences we used the estimated coefficients and SEs for these 220 probes generated in a seventh independent (Munich) cohort, which consisted of 450K data generated in the prefrontal cortex (N = 45) and sorted neuronal and non-neuronal nuclei from the occipital cortex (N = 26) (Table 1). This cohort had not been used in our discovery analyses as < 50 samples were available. Notably, we identified a similar pattern of Braak-associated DNA methylation changes for the 220 Bonferroni significant cross-cortex probes in this replication cohort, with a significantly correlated effect size between the discovery dataset and the replication prefrontal cortex (r = 0.64, P = 5.24 x 10^-27^), neuronal (r = 0.45, P = 1.56 x 10^-12^) and non-neuronal datasets (r = 0.79, P= 1.43 x 10^-47^) with a similar enrichment for the same direction of effect (sign test: prefrontal cortex P = 4.59 x 10^-28^, neuronal P = 6.13 x 10^-15^, non-neuronal P = 1.06 x 10^-42^) (Figure 3a). The most significant probe from the cross-cortex meta-analysis (cg12307200) showed consistent hypomethylation in disease in all cohorts in all cortical brain regions, with this direction of effect replicated in the prefrontal cortex and non-neuronal nuclei samples, but not the neuronal nuclei samples, suggesting that this is primarily driven by non-neuronal cell types, which are likely to be glia (Figure 3b). We have developed an online database (www.epigenomicslab.com/ad-meta-analysis/), which can generate a forest plot showing the ES and SE across any of the discovery cohorts and the Munich sample types for any of the 403,763 probes that passed our quality control. This allows researchers to determine the consistency of effects across cohorts for a given CpG site as well as the likely cellular origin of the signature. In addition, our tool can generate mini-Manhattan plots to show DMRs utilizing the summary statistics from the cross-cortex meta-analysis.

**Figure 3:**
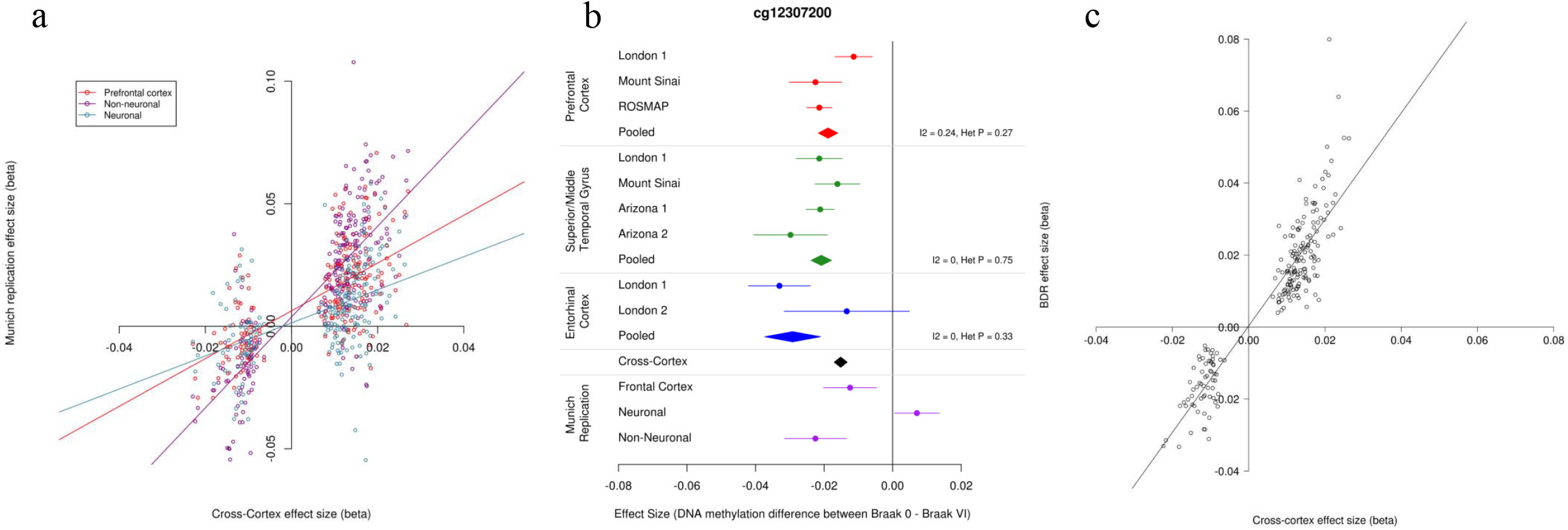
Independent replication of the Bonferroni significant cross-cortex differentially methylated loci. (**a**) The methylation (beta) effect size (ES) of the 220 cross-cortex differentially methylated positions (DMPs) identified in the discovery (N = 1,408) cohorts (X-axis) were significantly correlated with the ES in the Munich replication cohort in the prefrontal cortex (red; N = 45, r = 0.64, P = 5.24 x 10^-27^), sorted neuronal cells (light blue; N = 26, r = 0.45, P = 1.56 x 10^-12^) and non-neuronal cells (purple; r = 0.79, N = 26, P = 1.43 x 10^-47^) (Y-axis). (**b**) A forest plot of the most significant cross-cortex DMP (cg12307200, chr3:188664632, P = 4.48 x 10^-16^). The effect size is shown in the prefrontal cortex (red; N = 959), temporal gyrus (green; N = 608) and entorhinal cortex (blue; N = 189) for the different discovery cohorts. The X-axis shows the beta ES, with dots representing ES and arms indicating standard error (SE). ES from the intra-tissue meta-analysis using all available individual cohorts are represented by polygons in the corresponding tissue color. The black polygon represents the cross-cortex data. Shown in purple on the plot is the ES in the Munich replication cohort in the prefrontal cortex and sorted neuronal cells and non-neuronal cells, with the direction of effect suggesting the hypomethylation seen in the discovery cohorts is driven by changes in non-neuronal cells. (**c**) In the BDR replication cohort (N = 590) DNA methylation data was available in the prefrontal cortex for 208 of the 220 Bonferroni significant cross-cortex DMPs. The ES of these 208 cross-cortex DMPs in the discovery cohorts (X-axis) were significantly correlated with the ES in the BDR replication cohort (r = 0.53, P = 4.13 x 10^-16^) (Y-axis).

Finally, we had access to DNA methylation data generated in an eighth independent (Brains for Dementia Research [BDR]) cohort. This consisted of Illumina Infinium HumanMethylation EPIC BeadChip (EPIC array) data in the prefrontal cortex in 590 individuals^15^. As this is the successor to the 450K array (which had been used for the other seven cohorts), there are some differences in genome coverage, and for the 220 Bonferroni significant cross-cortex DMPs we had identified in the discovery cohorts, only 208 probes are also present on the EPIC array. For these overlapping 208 probes, we observed a significantly correlated effect size between the discovery dataset and the BDR dataset (r = 0.53, P = 4.13 x 10^-16^) (**Figure 3c**), with all 208 probes showing the same direction of effect (sign test P = 4.86 x 10^-63^).

### Cross-cortex AD-associated DMPs are enriched in specific genomic features

To identify if the cross-cortex DMPs reside in specific genomic features, we used a Fisher’s exact test to look for an enrichment of the 220 DMPs using Slieker annotations^16^ (Supplementary Data 11, Supplementary Figure 18). We observed a significant over representation of Bonferroni significant DMPs in CpG islands of gene bodies (odds ratio [OR] = 3.199, P = 4.76 x 10^-10^), and in CpG island shelves and non-CpG island areas of proximal promoters (OR = 3.571, P = 9.09 x 10^-3^ and OR = 1.641, P = 0.03, respectively). However, DMPs located in CpG islands in the proximal promoter were under-represented (OR = 0.353, P = 2.08 x 10^-6^). There was a significant over representation of the 220 cross-cortex DMPs in the first exon (OR = 1.80, P = 0.02), with an under enrichment within 1500bp of the transcription start site (OR = 0.49, P = 3.82 x 10^-3^) (Supplementary Data 12, Supplementary Figure 19).

### DNA methylomic signatures in the cortex can explain variance in the degree of pathology

We were interested to investigate whether the Braak-associated DNA methylation patterns we had identified across the cortex could accurately predict the pathological load of a brain sample and how much variance this explained. To this end we took samples within the discovery cohorts with either low pathology (Braak 0-II [controls]: N = 407), or high pathology (Braak V-VI [AD]: N = 589) and used these as a training dataset. We then used elastic net regression to identify 110 probes in the 220 cross-cortex Bonferroni significant loci (Supplementary Data 13) that were able to explain the most variance between post-mortem low pathology [control] from high pathology [AD] status in our training dataset (N = 996) (Supplementary Data 14, Figure 4). In our training data, we achieved an Area Under the Curve (AUC) of the Receiver Operating Characteristic (ROC) of 94.33% (CI = 92.88-95.64%, variance explained = 71.11%). We then tested its performance in the Munich replication samples (N = 38) and the BDR replication samples (N = 454), where it achieved an AUC of 73.95% (CI = 55.17-88.89%, variance explained = 20.18%) and 70.36% (CI = 65.52-75.12%, variance explained = 15.87%), respectively (Supplementary Data 14, Figure 4).

**Figure 4:**
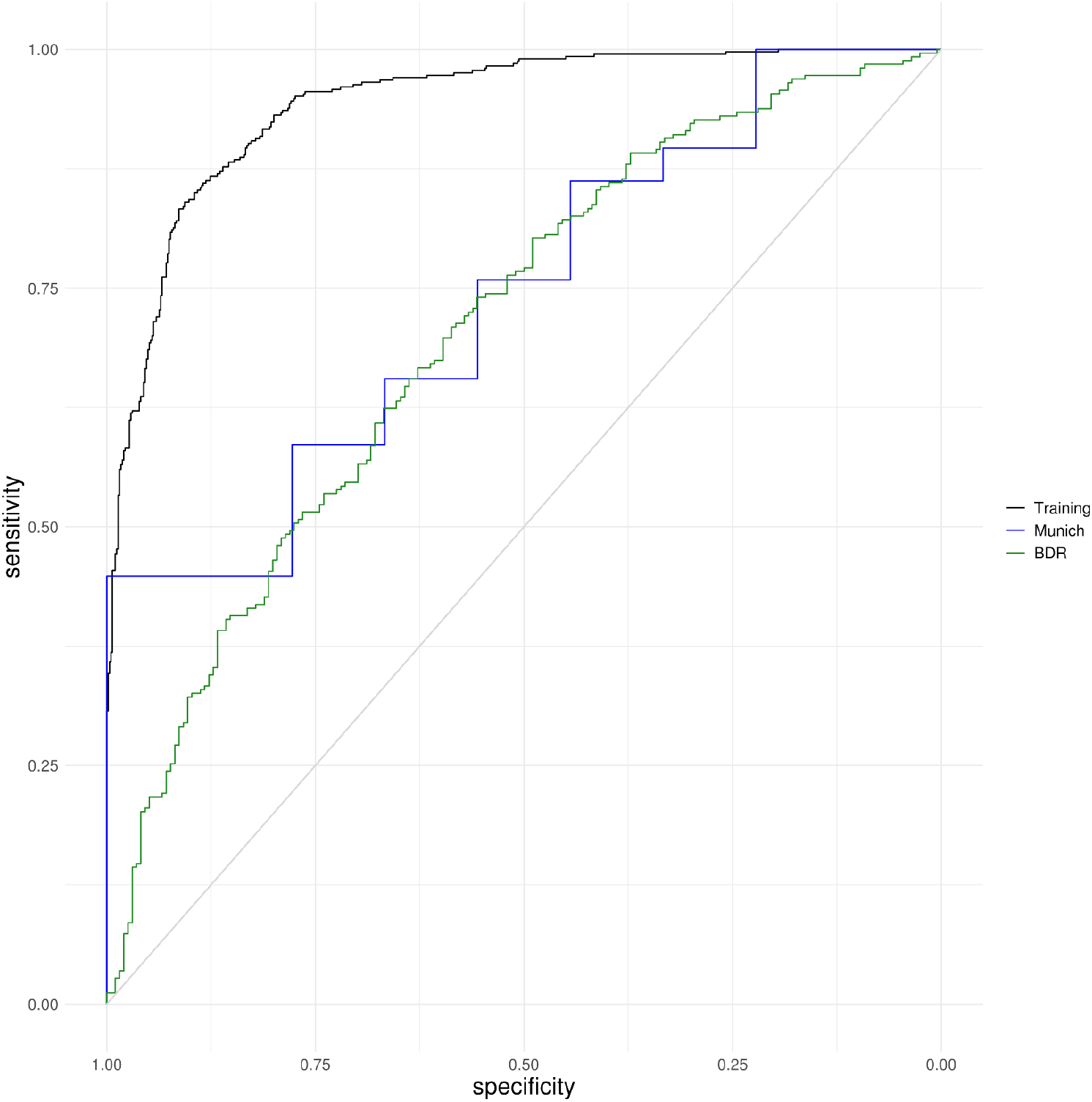
Receiver Operating Characteristic (ROC) graphs highlighting the Area Under the Curve (AUC) for the 110 cross-cortex probes that can best explain the variance in Braak pathology. An elastic net penalized regression model was used to identify a subset of 110 of the Bonferroni significant cross-cortex probes that could best predict whether a sample has low pathology (Braak 0-II: control) compared to high pathology (Braak V-VI: AD) in a training dataset comprised of 996 discovery samples (Braak 0-II: N = 407, Braak V-VI: N = 589). This model had an Area Under the ROC Curve (AUC) of 94.33% (confidence interval [CI] = 92.88-95.64%) and explained 71.11% of the pathological variance (black line). The 110 probe signature was then tested in two independent replication cohorts. In the Munich prefrontal cortex samples (Braak 0-II: N = 9, Braak V-VI: N = 29) the model had an AUC of 73.95% (CI = 55.17-88.89%), explaining 20.18% of the variance (blue line). In the BDR prefrontal cortex samples (Braak 0-II: N = 196, Braak V-VI: N = 258) the model had an AUC = 70.36% (CI = 65.52-75.12%), explaining 15.87% of the variance (green line). A list of the 110 probes and their performance characteristics can be found in Supplementary Data 13 and 14, respectively.

### DNA methylation signatures in AD cortex are largely independent of genetic effects

DNA methylomic variation can be driven by genetic variation via methylation quantitative trait loci (mQTLs). To explore whether SNPs may be driving the methylation differences we observed (in *cis)* we used the xQTL resource to identify *cis*-mQTLs associated with the 220 Bonferroni significant cross-cortex DMPs^17^. We identified 200 Bonferroni corrected mQTLs, which were associated with DNA methylation at 18 of the 220 cross-cortex DMPs (Supplementary Data 15). This suggests that the majority of Braak-associated DMPs are not the result of genetic variation in *cis*. None of these mQTLs overlapped with lead SNPs (or SNPs in LD) identified in the most recent genome-wide association study (GWAS) of diagnosed late-onset AD from Kunkle *et al*^18^. Next, we were interested in exploring whether DNA methylation is enriched in genes known to harbor AD-associated genomic risk variants. Using the AD variants from Kunkle *et al*^18^ we examined the enrichment of Braak-associated DNA methylation in 24 LD blocks harboring risk variants. Twenty of these LD blocks contained > 1 CpG site on the 450K array and using Brown’s method we combined P values within each of these 20 genomic regions. We observed Bonferroni-adjusted significant enrichment in the cross-cortex data in the *HLADRB1* (Chr6: 32395036-32636434: adjusted P = 1.20 x 10^-3^), *SPI1* (Chr11: 47372377-47466790, adjusted P = 5.76 x 10^-3^), *SORL1* (Chr11: 121433926-121461593, adjusted P = 0.019), *ABCA7* (Chr19: 1050130-1075979, adjusted P = 0.022) and *ADAM10* (Chr15: 58873555-59120077, adjusted P = 0.022) LD regions (Supplementary Data 16).

## DISCUSSION

This meta-analysis of AD EWAS utilizes six published independent sample cohorts with a range of cortical brain regions and cerebellum available as a discovery dataset. Two further independent cortical datasets where then used for replication, including data from sorted nuclei populations. Our data can be explored as part of an online searchable database, which can be found on our website (https://www.epigenomicslab.com/ad-meta-analysis). By performing a meta-analysis within each tissue, we have been able to identify 236, 95 and ten Bonferroni significant DMPs in the prefrontal cortex, temporal gyrus and entorhinal cortex, respectively. Although far fewer loci were identified in the entorhinal cortex compared to the other cortical regions, this is likely due to the reduced sample size in this tissue. In the cerebellum despite meta-analyzing > 500 unique samples, we identified no Braak-associated DNA methylation changes. Furthermore, there was no correlation of the effect size of Bonferroni significant DMPs identified in any of the cortical regions with the effect size of the same probes in the cerebellum. Taken together, this suggests that DNA methylomic changes in AD are cortex cell type specific. This observation is interesting as the cerebellum is said to be spared from AD pathology, with an absence of neurofibrillary tangles, although some diffuse amyloid-beta plaques are reported^19^. Interestingly, a recent spatial proteomics study of AD reported a large number of protein changes in the cerebellum in AD; however, the proteins identified were distinct from other regions examined and thus the authors suggested a potential protective role^20^.

Although many loci showed similar patterns of Braak-associated DNA methylation across the different cortical regions, some loci did show some regional specificity. In order to identify CpG sites that showed common DNA methylation changes in disease we performed a cross-cortex meta-analysis. Using this approach we identified 220 Bonferroni significant probes associated with Braak stage of which 46 probes had been previously reported at Bonferroni significance in the individual cohort studies that we had used for our meta-analysis, for example two probes in *ANK1*, four probes in *RHBDF2* and one probe in *HOXA3*, amongst others. Interestingly, our approach did identify 174 novel CpGs, corresponding to 102 unique genes, of which 84 genes had not been previously reported at Bonferroni significance in any of the previously published AD brain EWAS, highlighting the power of our meta-analysis approach for nominating new loci. This included 15 novel genes with at least two Bonferroni significant DMPs each, including five probes in *AGAP2*, three probes in *SLC44A2* and two probes each in *CDH9, CPEB4, DUSP27, GCNT2, MAMSTR, PTK6, RGMA, RHOB, SMURF1, THBS1, ZNF238* and *ZNF385A*. These genes had not been identified previously in an AD EWAS at this significance threshold, although a number of these genes had been previously identified from DMR analyses, which have a less stringent threshold. However, we did identify one novel gene (*HOXB3*) with three Bonferroni significant DMPs, which had not been identified at this significance threshold in previous EWAS DMP or DMR analyses in AD brain. The nomination of loci in the *HOXB* gene cluster is interesting; a recent study of human Huntington’s disease brain samples also highlighted significantly increased *HOXB3* gene expression in the prefrontal cortex^21^, an interesting observation given that both AD and Huntington’s disease are disorders that feature dementia. Furthermore, we have recently reported AD-associated hypermethylation of the *HOXB6* gene in AD blood samples^22^. Our pathway analysis highlighted differential methylation in a number of developmental pathways, mainly featuring the *HOXA* and *HOXB* gene clusters. Although it is unclear why developmental genes may be changed in a disease that primarily affects the elderly, it has been implied that genes such as these may be involved in neuroprotection after development^23^. A number of the other novel genes with multiple DMPs are also biologically relevant in the context of AD, for example *GCNT2* was recently shown to be differentially expressed in the Putamen between males and females with AD^24^. Interestingly, some of the protein products of genes we identified have also been previously linked with AD; PTK6 is a protein kinase whose activity has been shown to be altered in post-mortem AD brain^25^. Similarly, RGMA has been shown to be increased in AD brain, where it accumulated in amyloid-beta plaques^26^.

Our genomic enrichment analyses identified an over representation of hypermethylated loci in AD and methylation in specific genomic features, for example CpG islands in gene bodies, and shelves and non-CpG island regions in proximal promoters. We demonstrated that the majority of DMPs we identified (N = 202) were not driven by genetic variation as only 18 of the 220 CpG sites have reported mQTLs. However, we did observe a significant enrichment of cross-cortex loci in the LD regions surrounding the AD-associated genetic variants *HLADRB1, SPI1, SORL1, ABCA7* and *ADAM10* after controlling for multiple testing Finally, we have developed a classifier that could accurately predict control samples with low pathology, from those with a post-mortem AD diagnosis due to high pathology using methylation values for 110 of the 220 Bonferroni significant probes, further highlighting that distinct genomic loci reproducibly show epigenetic dysfunction in AD cortex. Although the clinical utility of such a classifier is limited as it is developed in post-mortem cortical brain tissue, it does illustrate that specific robust patterns of DNA methylation differences occur as the disease progresses. These signatures require further investigation as they could represent novel therapeutic targets, particularly given the classifier had an AUC > 70% in all the training and replication datasets. However, it is worth noting that the variance explained by the 110 CpG signature was lower in the replication datasets than the discovery samples, which could be due to a low sample number (Munich) or the different Illumina array platform (BDR).

There are some limitations with our study. First, as we have largely utilized methylation data generated in bulk tissue, this will contain a mixture of different cell types. Furthermore, it is known that the proportions of the major brain cell types are altered in AD, with reduced numbers of neurons and increased glia. As such, it is possible that the identified DNA methylation changes represent a change in cell proportions. To address this, we have included neuron/glia proportions as a co-variate in our models to minimize bias and used data from sorted cell populations as part of our replication. Although this is the optimal strategy for the current study given the EWAS data had already been generated, future EWAS should be undertaken on sorted cell populations with larger sample numbers than the Munich replication cohort, or ideally at the level of the single cell. It is important to note that the data from the sorted nuclei populations in the Munich replication cohort were generated in the occipital cortex, which was not a bulk tissue used for any of the discovery cohorts. In the future it would be interesting to explore whether different disease-associated DNA methylation signatures were observed in neurons and glia isolated from different cortical brain regions. Second, our study has utilized previously generated EWAS data generated on the 450K array or EPIC array. Although the Illumina array platform has been the most widely used platform for EWAS to date, it is limited to only analyzing a relatively small proportion of the potential methylation sites in the genome (~400,000 on the 450K array) and given the falling cost of sequencing, future studies could exploit this by performing reduced representation bisulfite sequencing to substantially increase the coverage. In our study we have primarily used the UCSC annotation provided by Illumina to identify the gene relating to each DMP. However, this can lead to the annotation of overlapping genes, or no gene annotation, which can make it difficult to establish the gene of interest in the absence of functional studies. Our study has primarily focused on the results of a fixed-effects meta-analysis, as the majority of Bonferroni-significant DMPs do not display a high degree of heterogeneity. However, ~15% of the cross-cortex DMPs did have a significant heterogeneity P value and in this instance, it is worthwhile also considering the results of the random-effects meta-analysis. Although this heterogeneity could be driven by differences between cohorts, it is also plausible that it may be driven by tissue-specific effects as we used different cortical brain regions in the model. For example cg22962123 annotated to the *HOXA3* gene has a significant heterogeneity P value in the cross-cortex meta-analysis, but we had already shown this loci to be differentially methylated in the prefrontal cortex and temporal gyrus, but not the entorhinal cortex in our intra-tissue meta-analysis.

Another limitation of our study is that we have focused our analyses on Braak (neurofibrillary tangle)-associated methylation changes, as this measure was available in all cohorts. Given that amyloid-beta is another neuropathological hallmark of AD, it would also be of interest to identify neuritic plaque-associated DMPs. Unfortunately, this was not feasible in the current study as this measure was not available in all samples. In a similar vein, we did not exclude individuals with mixed pathology, or protein hallmarks of other neurodegenerative diseases, such as the presence of lewy bodies, or TDP-43 pathology. In the future, larger meta-analyses should stratify by the presence of these protein aggregates, particularly given that very few EWAS have been undertaken in other dementias. Indeed, only three DNA methylomic studies have been undertaken in cortical samples of individuals with other dementias to date^27–30^, with none of these studies utilizing > 15 individuals for EWAS. Further studies exploring common and unique DNA methylation signatures and our classifier in other diseases characterized by dementia will be vital for identifying disease-specific epigenetic signatures that could represent novel therapeutic targets. Finally, one key issue for epigenetic studies in post-mortem tissue is the issue of causality, where it is not possible to determine whether disease-associated epigenetic loci are driving disease pathogenesis, or are a consequence of the disease, or even the medication used for treatment. One method that can be used to address this is Mendelian Randomization^31^ however, this does require the CpG site to have a strong association with a SNP. Given that we only identified mQTLs at 18 of the 220 Bonferroni significant cross-cortex DMPs, this approach is not suitable for most of the loci we identified. At an experimental level establishing causality is difficult to address in post-mortem human studies, and therefore longitudinal studies in animal models, or modelling methylomic dysfunction through epigenetic editing *in vitro* will be useful approaches to address these issues. In addition, examining DNA methylation signatures in brain samples in pre-clinical individuals (*i.e*. during midlife) will be important for establishing the temporal pattern of epigenetic changes relative to the pathology.

In summary we present intra-tissue and cross-cortex meta-analyses of AD EWAS, highlighting numerous Bonferroni significant DMPs in the individual cortical regions and across the cortex, but not in the cerebellum, which were replicated in two independent cohorts. A number of these loci are novel and warrant further study to explore their role in disease etiology. We highlight that the nominated epigenetic changes are largely independent of genetic effects, with only 18 of the 220 Bonferroni significant DMPs showing a mQTL. We provide evidence that robust epigenomic changes in the cortex can predict the level of pathology in a sample. Looking to the future it will be important to explore the relationship between DNA methylation and gene expression in AD brain.

## METHODS

### Cohorts

Six sample cohorts were used for discovery in this study as they all had DNA methylation data generated on the 450K array for > 50 donors, enabling us to take a powerful meta-analysis approach to identify DNA methylation differences in AD. As our analyses focused specifically on neuropathology (tau)-associated differential methylation, inclusion criteria for all samples used in the discovery or replication cohorts was having post-mortem neurofibrillary tangle Braak stage available. For each discovery sample cohort DNA methylation was quantified using the 450K array. The London 1 cohort comprised of prefrontal cortex, superior temporal gyrus, entorhinal cortex, and cerebellum tissue obtained from 113 individuals archived in the MRC London Neurodegenerative Disease Brain Bank and published by Lunnon *et al*.^5^. The London 2 cohort comprised entorhinal cortex and cerebellum samples obtained from an additional 95 individuals from the MRC London Neurodegenerative Disease Brain Bank published by Smith and colleagues^8^. The Mount Sinai cohort comprised of prefrontal cortex and superior temporal gyrus tissue obtained from 146 individuals archived in the Mount Sinai Alzheimer’s Disease and Schizophrenia Brain Bank published by Smith and colleagues^7^. The Arizona 1 cohort consisted of 302 middle temporal gyrus and cerebellum samples from The Sun Health Research Institute Brain Donation Program^32^ published by Brokaw *et al*.^12^. The Arizona 2 cohort consisted of an additional 88 temporal gyrus and cerebellum samples from Lardonije *et al*.^10^. The ROSMAP cohort consisted of 709 samples from the Rush University Medical Center: Religious Order Study (ROS) and the Memory and Aging Project (MAP), which were previously published by De Jager and colleagues^6^. For replication purposes we used two further replication datasets. The Munich cohort from Neurobiobank Munich (NBM), which had bulk prefrontal cortex 450K array data from 45 donors, and 450K array data from fluorescence-activated cell sorted neuronal and non-neuronal (glial) populations from the occipital cortex from 26 donors as described by Gasparoni *et al*.^11^. The Brains for Dementia Research (BDR) cohort consisted of bulk prefrontal cortex Illumina Infinium EPIC array data from 590 donors, as described by Shireby *et al*^15^. Demographic information for all eight cohorts is available in Table 1. Ethical approval for the study was granted from the University of Exeter Medical School Research Ethics Committee (13/02/009).

### Data quality control and harmonization

All computations and statistical analyses were performed using R 3.5.2^33^ and Bioconductor 3.8^34^. A *MethylumiSet* object was created from iDATs using the methylumi package^35^ and *RGChannelSet* object was created using the minfi package^36^. Samples were excluded from further steps if (a) the mean background intensity of negative probes < 1,000, (b) the mean detection P values > 0.005, (c) the mean intensity of methylated or unmethylated signals were three standard deviations above or below the mean, (d) the bisulfite conversion efficiency < 80%, (e) there was a mismatch between reported and predicted sex, or (f) the 65 SNP probes on the array show a modest level of correlation (using a cut-off of 0.65) between two samples (whereby the sample with the higher Braak score was retained). Sample and probe exclusion was performed using the *pfilter* function within the wateRmelon package^37^, with the following criteria used for exclusion: samples with a detection P > 0.05 in more than 5% of probes, probes with < three beadcount in 5% of samples and probes having 1% of samples with a detection P value > 0.05. Finally, probes with common (minor allele frequency > 5%) SNPs in the single base extension position or probes that are nonspecific or mis-mapped were excluded^38,39^, leaving 403,763 probes for analysis. Samples numbers after quality control are those shown in Table 1.

Quantile normalization was applied using the *dasen* function in the wateRmelon package^37^. For the discovery cohorts, DNA methylation data was corrected by regressing out the effects of age and sex in all samples in each cohort and tissue separately, with neuron/glia proportions included as an additional covariate in cortical regions. The neuron/glia proportions were calculated using the CETS package^40^, and were not included as a co-variate for the cerebellum as the neuronal nuclear protein (NeuN) that was used to generate the neuron/glia algorithm is not expressed by some cerebellar neurons^41^. These three variables (age, sex, neuron/glia proportions) were regressed out of the data as we found that they strongly correlated with either of the first two principal components of the DNA methylation data in most of the datasets. Other potential sources of technical and biological variation (post-mortem interval, ancestry, plate, chip, study and bisulfite treatment batch) did not correlate as strongly with methylation in most datasets. We opted to use surrogate variables as a consistent method to control for variation derived from these measured and other unknown variables across all datasets. Surrogate variables were calculated using the *sva* function in the SVA package^42^. Linear regression analyses were then performed with respect to Braak stage (modelled as a continuous variable) using residuals and a variable number of surrogate variables for each study until the inflation index (lambda) fell below 1.2 (see Supplementary Data 17). The surrogate variables included for each cohort correlated with the technical and biological variables that we had not regressed out earlier, demonstrating that this method appropriately controlled for variation not driven by Braak stage. Quantile-quantile plots for the four intra-tissue and the cross-cortex meta-analyses are shown in Supplementary Figure 20. Although it appears from these plots that there is P value inflation, it is worth noting that (a) lambda for all meta-analyses < 1.2 and (b) P value inflation is commonly observed in many DNA methylation studies and standard methods to control for this in GWAS are not suitable for EWAS data^43^.

### Intra-tissue meta-analysis

We used the estimated coefficients and SEs from the six discovery cohorts to undertake an inverse variance intra-tissue meta-analysis independently in each available tissue using the *metagen* function within the Meta package^44^, which applies inverse variance weighting. The estimates and SEs from individual cohort Braak linear regression analyses were added to the model for each tissue. The prefrontal cortex analyses included three cohorts (N = 959: London 1, Mount Sinai, ROSMAP), the temporal gyrus analyses included four cohorts (N = 608: London 1, Mount Sinai, Arizona 1, Arizona 2) and the entorhinal cortex analyses included two cohorts (N = 189: London 1, London 2). The cerebellum analyses included data from four cohorts (N = 533: London 1, London 2, Arizona 1 and Arizona 2) although the cerebellum data for the Arizona 1 and 2 cohorts was generated in the same experiment, and so these were combined together as a single dataset. The ESs and corresponding SEs reported in this study correspond to the corrected DNA methylation (beta) difference between Braak 0 and Braak VI individuals. Bonferroni significance was defined as P < 1.238 x 10^-7^ to account for 403,763 tests. A fixed effects meta-analysis are the results primarily reported as it is the most appropriate model for our study as it can more reliably estimate the pooled effect and therefore the standard error and P value. However, in the Supplementary Data we do also report the results of the random effects meta-analysis as ~10% of Bonferroni significant DMPs in the intra-tissue meta-analysis had high heterogeneity and in which case the results from the random-effects model should also be considered.

### Cross-cortex meta-analysis

As multiple cortical brain regions were available for the London 1 and Mount Sinai cohorts, a mixed model was performed using the *lme* function within the nlme package^45^. Estimate coefficients and SEs from each EWAS were extracted and were subjected to bacon^43^ to control for bias and inflation, after which a fixed-effect inverse variance meta-analysis was performed across all discovery cohorts using the *metagen* function. A fixed effects model was selected in this instance for consistency with the intra-tissue meta-analysis, although the random effects meta-analysis results also shown in Supplementary Data 7.

### Replication analyses

For the Munich replication cohort, we extracted the beta values for the 220 cross-cortex Bonferroni significant DMPs. This DNA methylation data was then corrected for age, sex and neuron/glia proportions (bulk tissue only) prior to performing a linear regression analysis with respect to Braak stage. For the BDR replication cohort, we were provided with beta values for the 208 cross-cortex Bonferroni significant DMPs that were present on the EPIC array. This data had been corrected for age, sex, neuron/glia proportions, batch and principal component 1, before the linear regression analysis was performed with respect to Braak stage, with Bacon used to control for inflation. Additional information on the BDR dataset can be found in Shireby *et al*^15^.

### Annotations, pathway and regional analyses

Probes were annotated for tables using both the Illumina (UCSC) gene annotation (which is derived from the genomic overlap of probes with RefSeq genes or up to 1500bp from the transcription start site of a gene) and Genomic Regions Enrichment of Annotations Tool (GREAT)^46^ annotation version 4.0.4 (which annotates a DMP to genes with a transcription start site within 5kb upstream, or 1kb downstream). Pathway analyses were performed on the Illumina (UCSC) annotated genes corresponding to the 220 Bonferroni significant cross-cortex DMPs (N = 121 genes). We used the ‘gometh’ function within the missMethyl package (version 1.20.0)^47^, which performs one-sided hypergeometric tests and adjusts the test for the uneven number of probes per gene and pathway redundancy. The identified GO terms were subjected to the online tool REViGO (available at http://revigo.irb.hr/)^48^, to reduce the number of redundant functional terms based on semantic similarity between ontology terms. Resnik’s measure was used to compute the similarity of terms and a medium between terms similarity of 0.7 was allowed. As methylation at neighboring CpG sites can be highly correlated we used a method developed to identify SNPs in LD to identify independent signals^14^. For the 220 Bonferroni significant cross-cortex DMPs we used a threshold of r < 0.6 over 1mb to identify 165 independent (non-highly correlated) methylation signals. To identify DMRs consisting of multiple DMPs we used the Python package comb-p^49^ with a distance of 500bp and a seeded P value of 1.0 x 10^-4^. Comb-p was selected for DMR identification over alternative methods as it uses P values as input and so was the most suitable method for calling DMRs in the cross-cortex meta-analysis where multiple brain regions were available for some of the individuals. We have used comb-p to call DMRs in a number of our previous EWAS in AD, including studies where we have validated the top DMRs using an alternative technology such as pyrosequencing^5,8,22^.

### Genomic enrichment analyses

To test for an enrichment of DMPs in specific genomic features (*i.e*. CpG islands, shelves, shores, non-CpG island regions) in certain genomic regions (*i.e*. intergenic, distal promoter, proximal promoter, gene body, downstream) we annotated all DMPs with Slieker annotation^16^ and performed a two-sided Fisher’s exact test comparing to all probes analyzed (N = 403,763). We also used a Fisher’s exact test to test for an enrichment of DMPs in genomic regions related to transcription based on the Illumina annotation (TSS1500, TSS200, 5’ UTR, 1^st^ exon, gene body, 3’ UTR). To investigate whether any of the 220 Bonferroni significant cross-cortex DMPs were driven by genetic variation we used the xQTL resource to identify which of these DMPs are established *cis*-mQTLs^17^. To explore whether Braak-associated methylation was enriched in known AD GWAS variants we used Brown’s method to combine together P values from our meta-analyses for probes residing in the LD blocks around the genome-wide significant (P < 5.0× 10^-8^) GWAS variants identified by the stage one meta-analysis of Kunkle *et al*.^18^ Of the 24 LD blocks reported by Kunkle and colleagues, 20 contained > 1 CpG site on the 450K array and the P values for each CpG in a given block were combined using Brown’s method, which accounts for the correlation structure between probes, with the regional P values adjusted to correct for multiple testing.

### Quantifying variance in Braak pathology explained by DNA methylation signatures

For this analysis control samples (Braak low [0-II]: N = 407) and AD cases (Braak high [VVI]: N = 589) from the cross cortex discovery dataset were used for training a classifier. A penalized regression model was used to select the optimum (N = 110) CpG probes from the 220 cross-cortex Bonferroni significant DMPs that determined case-control status in the training dataset using the R package GLMnet^50^. Elastic net uses a combination of ridge and lasso regression, in which alpha (α) = 0 corresponds to ridge, whilst α = 1 corresponds to lasso, the elastic net α parameter used was 0.5. The lambda value was derived when using 10-fold cross validation on the training dataset. The model was then tested for AUC ROC value, confidence intervals (CI) and variance explained in the testing dataset as well as the independent replication Munich (Braak 0-II: N = 9, Braak V-VI: N = 29) and BDR (Braak 0II: N = 196, Braak V-VI: N = 258) prefrontal cortex datasets.

## Data availability

The data supporting the findings of this study are available within the article, Supplementary Information or from the authors upon request. Some of the datasets are also available on GEO including London 1 data (GSE59685 [https://www.ncbi.nlm.nih.gov/geo/query/acc.cgi?acc=GSE59685]), London 2 data (GSE105109 [https://www.ncbi.nlm.nih.gov/geo/query/acc.cgi?acc=GSE105109]), Mount Sinai data (GSE80970 [https://www.ncbi.nlm.nih.gov/geo/query/acc.cgi?acc=GSE80970]), Arizona 1 data (GSE134379 [https://www.ncbi.nlm.nih.gov/geo/query/acc.cgi?acc=GSE134379]), Arizona 2 data (GSE109627 [https://www.ncbi.nlm.nih.gov/geo/query/acc.cgi?acc=GSE109627]) and Munich data (GSE66351 [https://www.ncbi.nlm.nih.gov/geo/query/acc.cgi?acc=GSE66351]). The BDR data is available from the authors upon reasonable request. We have developed an online database, which can present summary statistics, which is available from our website: www.epigenomicslab.com/ad-meta-analysis/. All scripts for data analyses performed in this manuscript can be found at: https://github.com/rgs212/Meta-analysis-Smith^51^.

## Acknowledgements

This work was funded by a major project grant from the Alzheimer’s Society UK (AS-PG-14-038) to KL, an Alzheimer’s Association US New Investigator Research Grant (NIRG-14-320878) to KL and a project grant from the Medical Research Council (MRC) (MR/N027973/1) to KL as part of a larger collaborative project funded to KL and DLAvdH for the EPI-AD consortium through the Joint Programme—Neurodegenerative Disease Research (JPND) initiative. Data analysis was undertaken using high-performance computing supported by a Medical Research Council (MRC) Clinical Infrastructure Award (M008924) to JM. The project was also supported through PhD studentships from the Alzheimer’s Society (GS), BRACE (Bristol Research into Alzheimer’s and Care of the Elderly) (GW) and the MRC GW4 Doctoral Training Program (DTP) (JAYR). BDR is jointly funded by Alzheimer’s Research UK and the Alzheimer’s Society in association with the MRC. DNA methylation data generated in the Brains for Dementia Research cohort was supported by the Alzheimer’s Society and Alzheimer’s Research UK (ARUK). Research reported in this publication was also supported by the National Institute on Aging of the National Institutes of Health under award numbers P30AG19610, P30AG10161, R01AG15819, R01AG17917, R01AG36042, R01AG036039, R01AG036400 and R01AG067015. The content is solely the responsibility of the authors and does not necessarily represent the official views of the National Institutes of Health. We thank the donors and families who have made this research possible.

## Contributions

ARS and GW conducted laboratory experiments. RGS, EP, GS, EH, WV and MW undertook data analysis, bioinformatics and/or support with data review. RGS, EP, ARS, JAYR, DM, GG, MR, AG, AJS, LS, VH, DLAvdH, DB, PTF, AJT, SL, KM, JW, PDC, DAB, PLDJ, JM and KL provided data for the meta-analysis. LS developed the online database. KL conceived of the idea and directed the project. KL, RGS and EP drafted the manuscript. All authors read and approved the final submission.

