## Supplementary Information (Supplementary Figures) for "A meta-analysis of epigenome-wide association studies in Alzheimer’s disease highlights novel differentially methylated loci across cortex"

**Supplementary Figure 1: Workflow for meta-analyses.** Raw iDATs for each tissue and cohort were loaded into R as a MethylumiSet and RGChannelSet before stringent quality control. Data was then harmonized via quantile normalization and regressing out the co-variates of age, sex and cell proportions (proportions used in cortex bulk data only). Subsequently, individual epigenome-wide association studies (EWAS) were performed in each tissue and cohort with respect to Braak stage. Next, we used the effect size (ES) and standard error (SE) from the individual EWAS within a given tissue in an inverse variance fixed-effect meta-analysis. For the cross-cortex meta-analysis we first used a mixed effects model with nested IDs in donors with multiple tissues from the same cohort. We then performed an inverse variance fixed-effect meta-analysis using these ES and SE and those from samples with just a single brain region. Downstream analyses on the outputs of this meta-analysis included pathway analyses, genomic enrichment analyses and classification analyses.

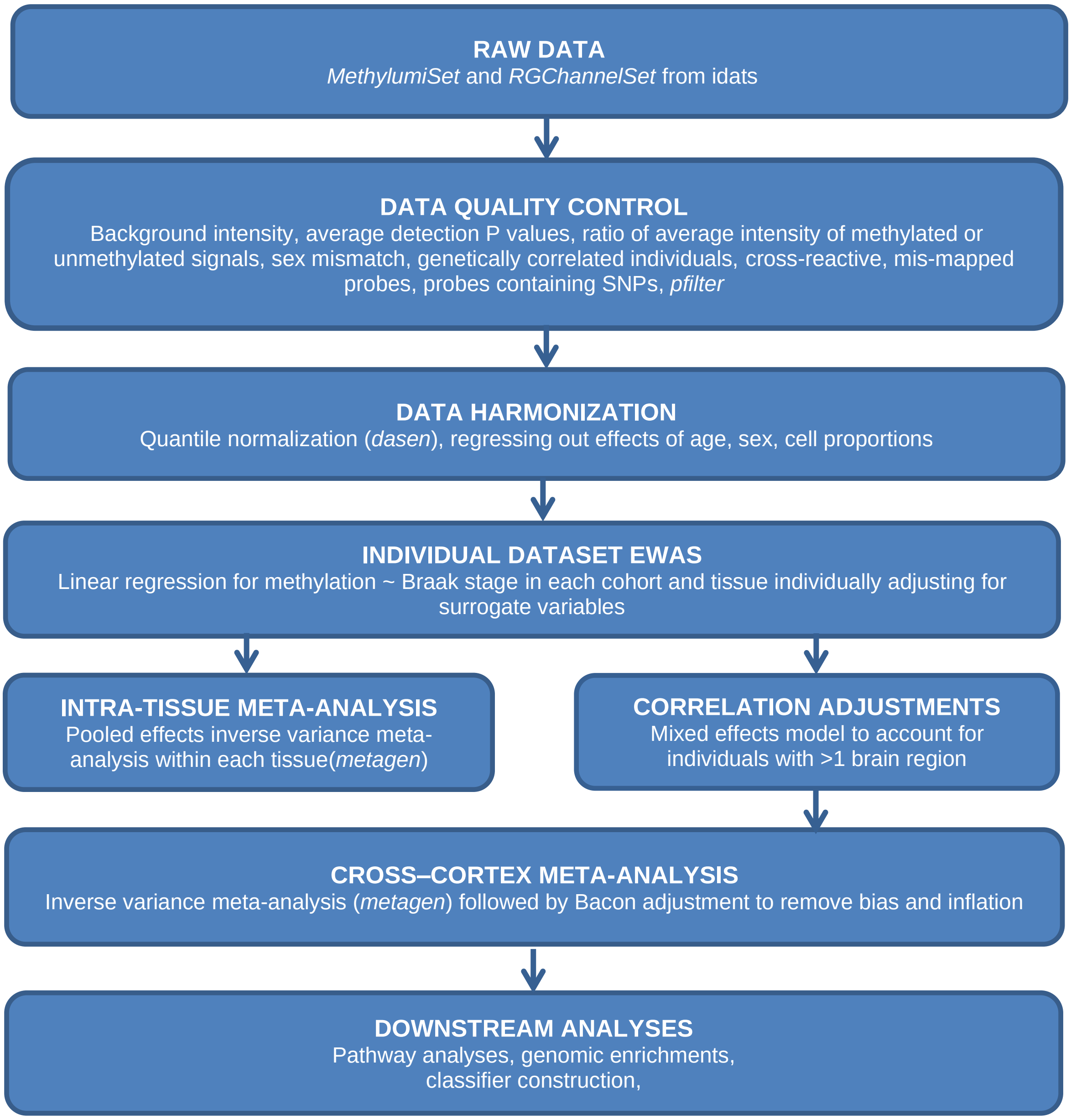

**Supplementary Figure 2: Volcano plot of differentially methylated positions (DMPs) identified in the prefrontal cortex inverse variance fixed effects meta-analysis (N =959)**. The X-axis shows methylation (beta) effect size (ES) and the Y-axis shows -log-10(p). Gray probes indicate an ES ≥ 0.01, whilst the blue probes indicate an ES ≥ 0.01 and a Bonferroni significant P-value ($P < 1.238 \times 10^{-7}$). Exact p-values are provided in Supplementary Data 1.

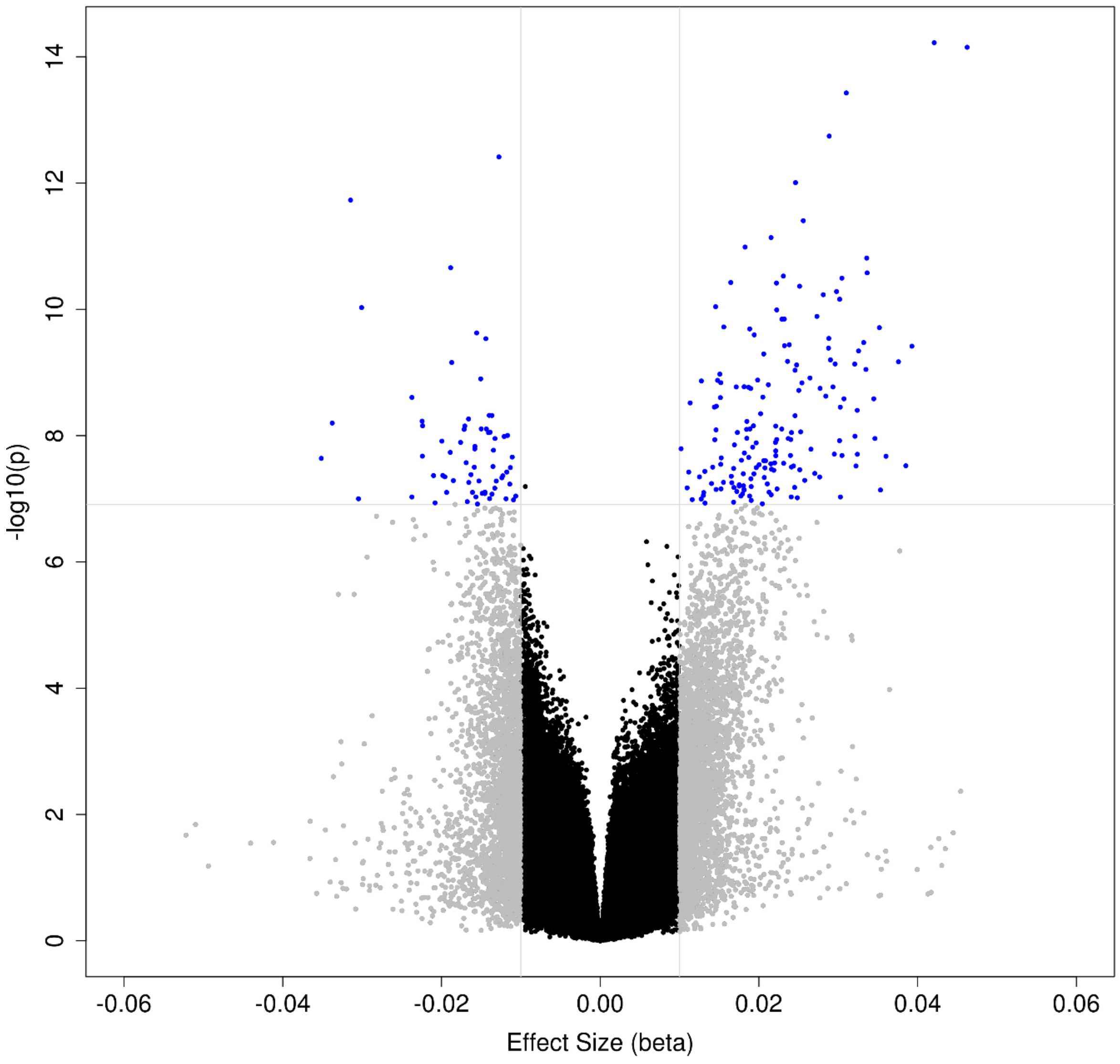

**Supplementary Figure 3: Forest plot of the most significant differentially methylated position (DMP) in the prefrontal cortex inverse variance fixed effects meta-analysis (cg22962123).** The methylation (beta) effect size (ES) is shown in the prefrontal cortex (red; N = 959), temporal gyrus (green; N = 608) and entorhinal cortex (blue; N = 189) for the different cohorts. The X-axis shows the beta ES, with dots representing ES and arms indicating standard error (SE). ES from the intra-tissue meta-analysis using all available individual cohorts are represented by polygons.

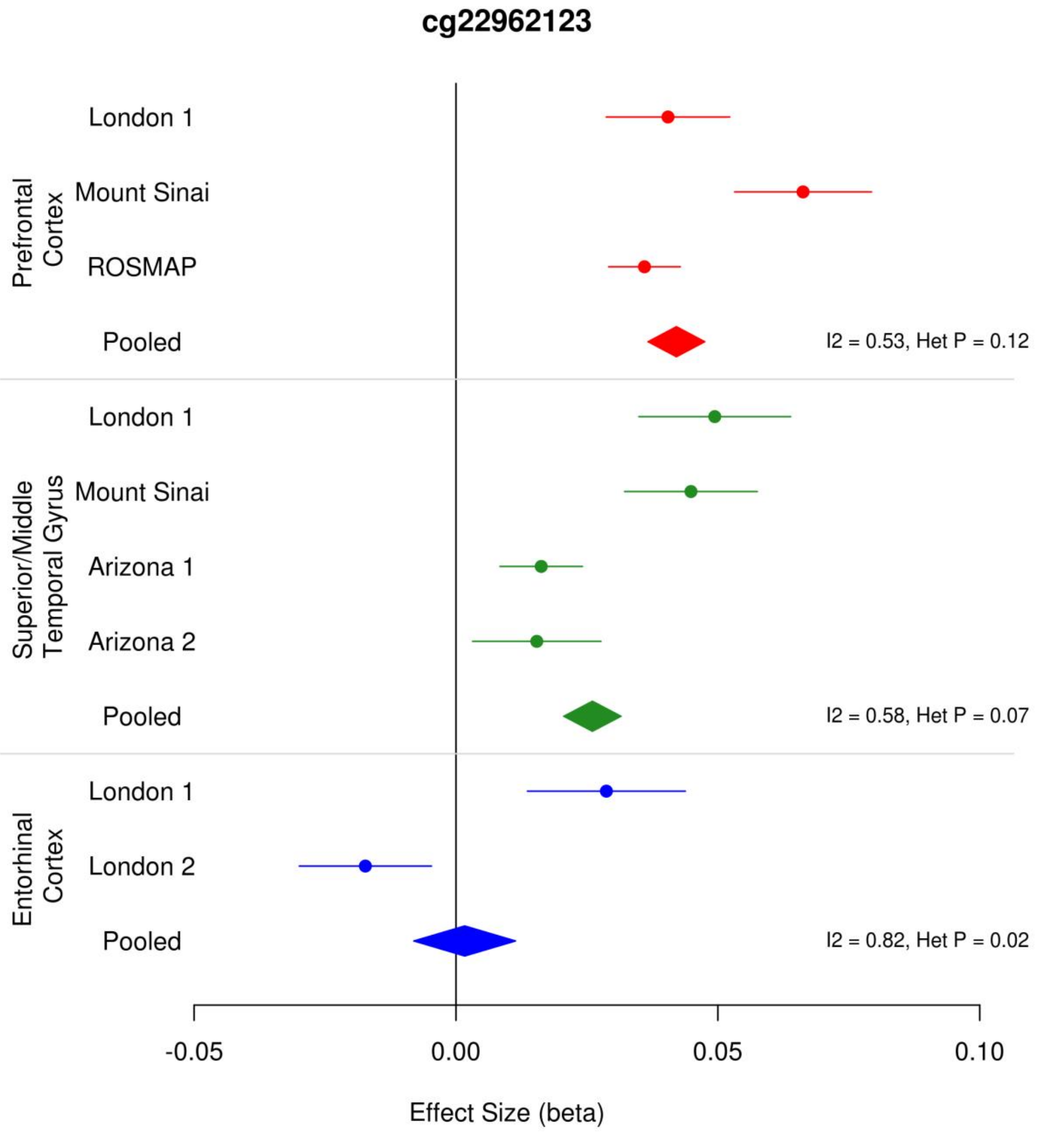

**Supplementary Figure 4: Neuropathology-associated Bonferroni significant differentially methylated positions (DMPs) identified in the prefrontal cortex meta-analysis are replicated in other cortical regions.** The methylation (beta) effect size (ES) of the 236 Braak-associated Bonferroni significant DMPs identified in the prefrontal cortex (N = 959) from our inverse variance fixed effect meta-analysis model (X-axis) were significantly correlated with the ES of the same probes in the temporal gyrus (green; N = 608, r = 0.94, P = 6.17 x $10^{-112}$) and entorhinal cortex (blue; N = 189, r = 0.58, P = 1.80 x $10^{-22}$) (Y-axis).

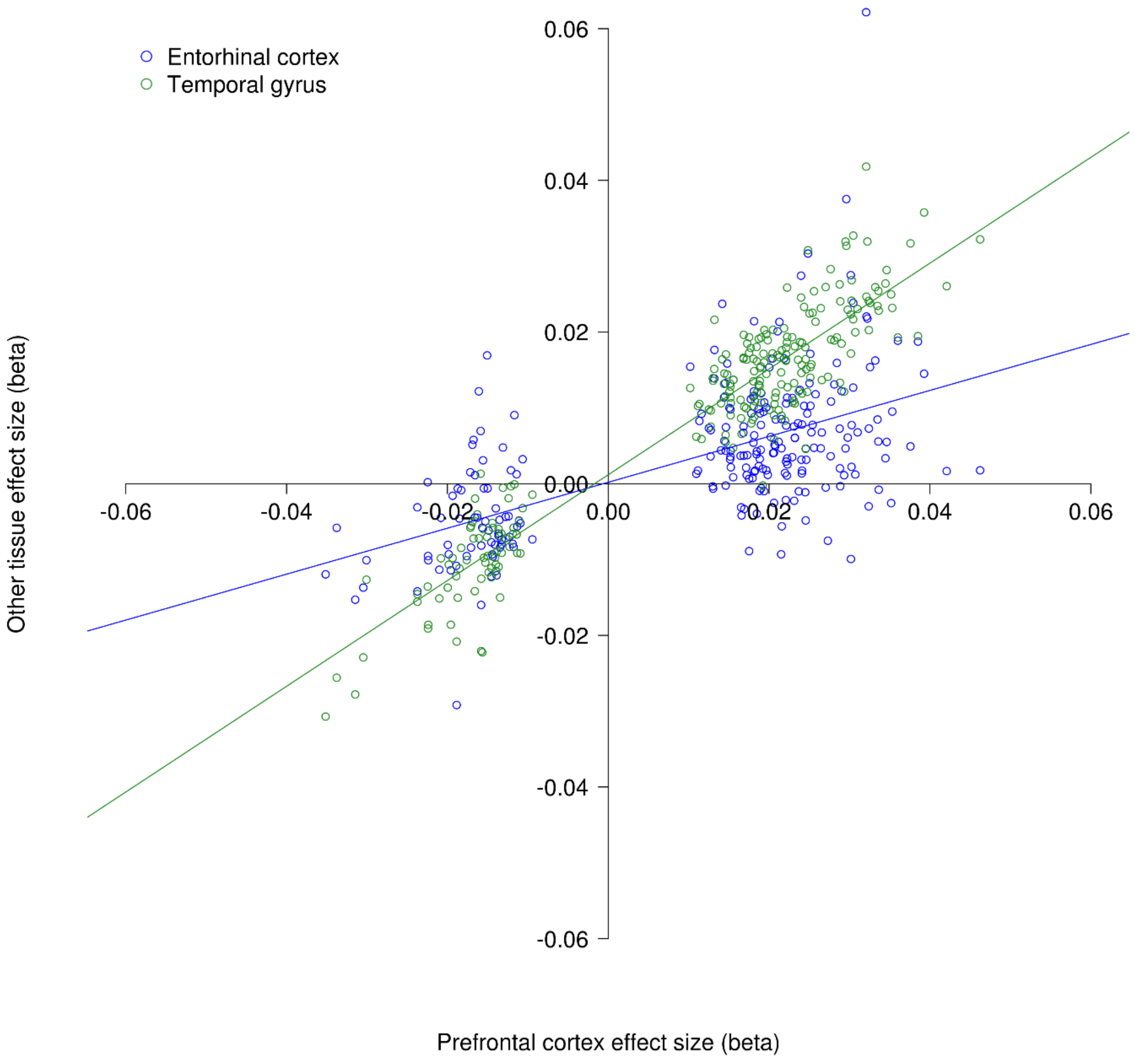

**Supplementary Figure 5: The most significant differentially methylated region (DMR) in the prefrontal cortex inverse variance fixed effects meta-analysis (chr7:26955524-27473741) contained 20 probes and resided in the *HOXA* region (N = 959).** The horizontal red line denotes the Bonferroni significance level of $P < 1.238 \times 10^{-7}$. Red probes represent a positive methylation (beta) effect size (ES) $\geq$ 0.01, blue probes represent a negative ES $\geq$ 0.01. Underneath the gene tracks are shown in black with CpG islands in green.

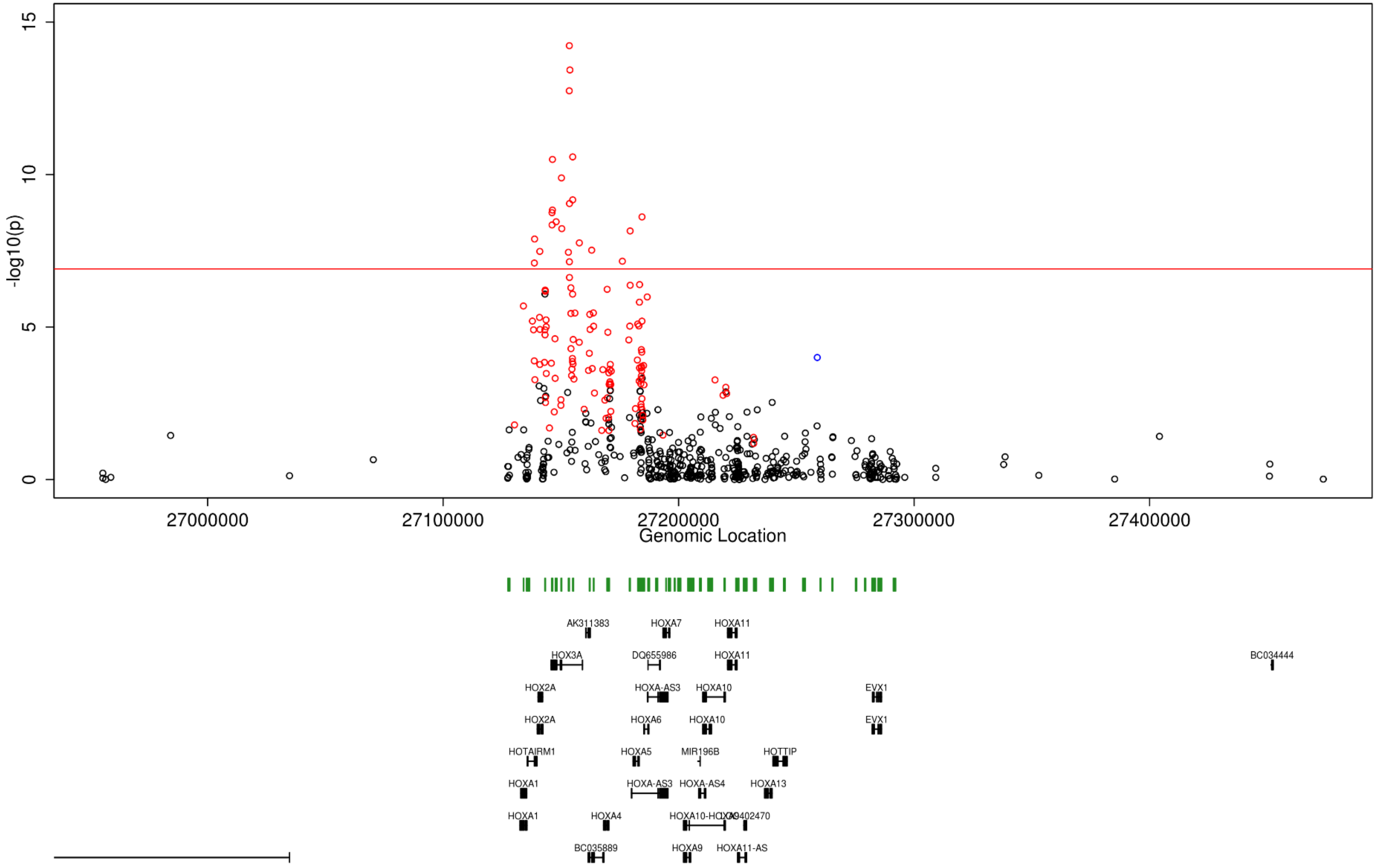

**Supplementary Figure 6: Volcano plot of differentially methylated positions (DMPs) identified in the temporal gyrus inverse variance fixed effects meta-analysis (N =608).** The X-axis shows beta effect size (ES) and the Y-axis shows -log10(p). Gray probes indicate an ES ≥ 0.01, whilst blue probes indicate an ES ≥ 0.01 and a Bonferroni significant P-value ($P < 1.238 \times 10^{-7}$). Exact p-values are provided in Supplementary Data 3.

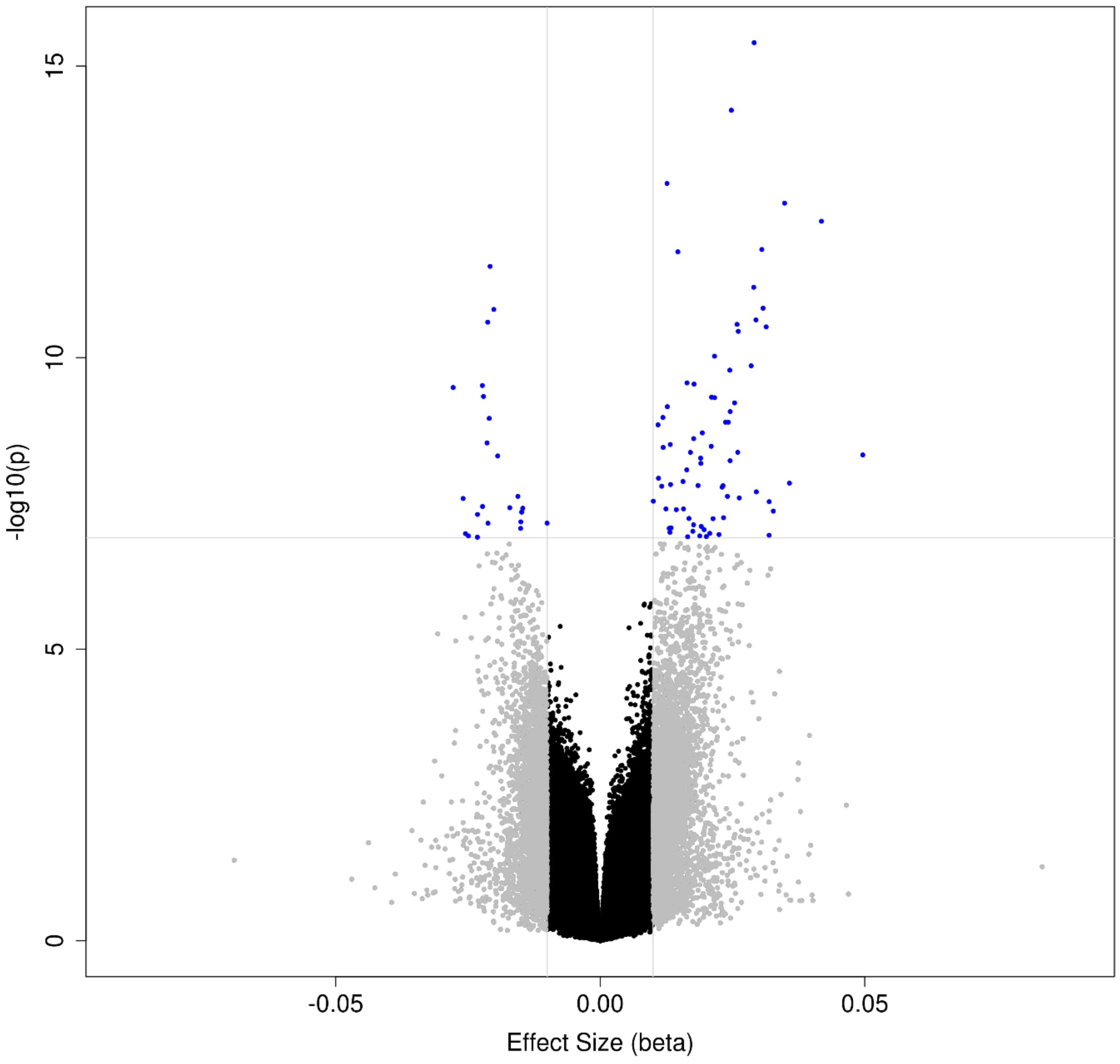

**Supplementary Figure 7: Forest plot of the most significant differentially methylated position (DMP) in the temporal gyrus and entorhinal cortex inverse variance fixed effects meta-analysis (cg11823178).** The methylation (beta) effect size (ES) is shown in the prefrontal cortex (red; N = 959), temporal gyrus (green; N = 608) and EC entorhinal cortex (blue; N = 189) for the different cohorts. The X-axis shows the beta ES, with dots representing ES and arms indicating standard error (SE). ES from the intra-tissue meta-analysis using all available individual cohorts are represented by polygons.

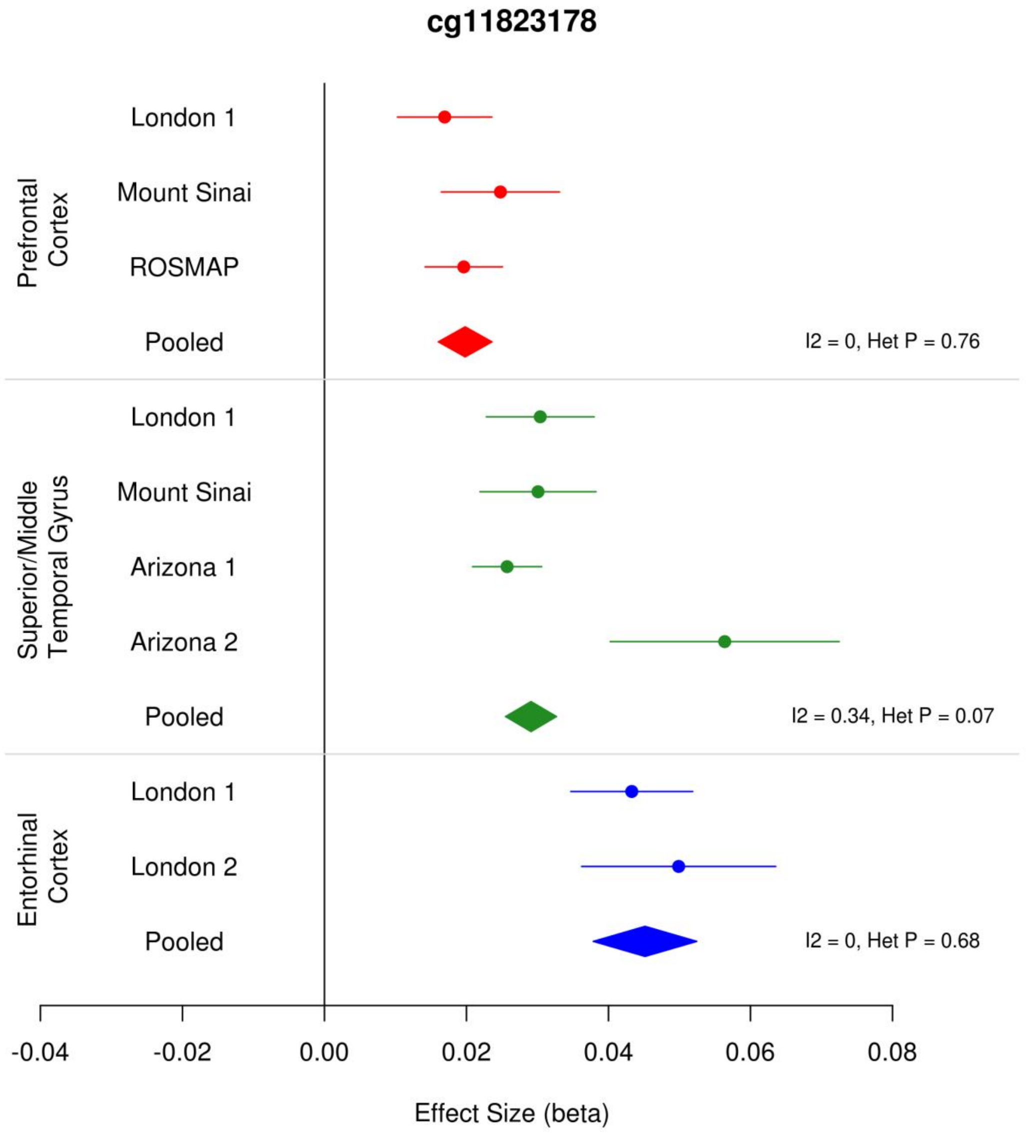

**Supplementary Figure 8: Neuropathology-associated Bonferroni significant differentially methylated positions (DMPs) identified in the temporal gyrus meta-analysis are replicated in other cortical regions.** The methylation (beta) effect size (ES) of Braak-associated Bonferroni significant DMPs identified in the temporal gyrus (N = 608) from our inverse variance fixed effect meta-analysis model (X-axis) were significantly correlated with the ES of the same probes in the prefrontal cortex (red; N = 959, r = 0.91, P = 5.09 x $10^{-38}$) and entorhinal cortex (blue; N = 189, r = 0.77, P = 4.02 x $10^{-20}$) (Y-axis).

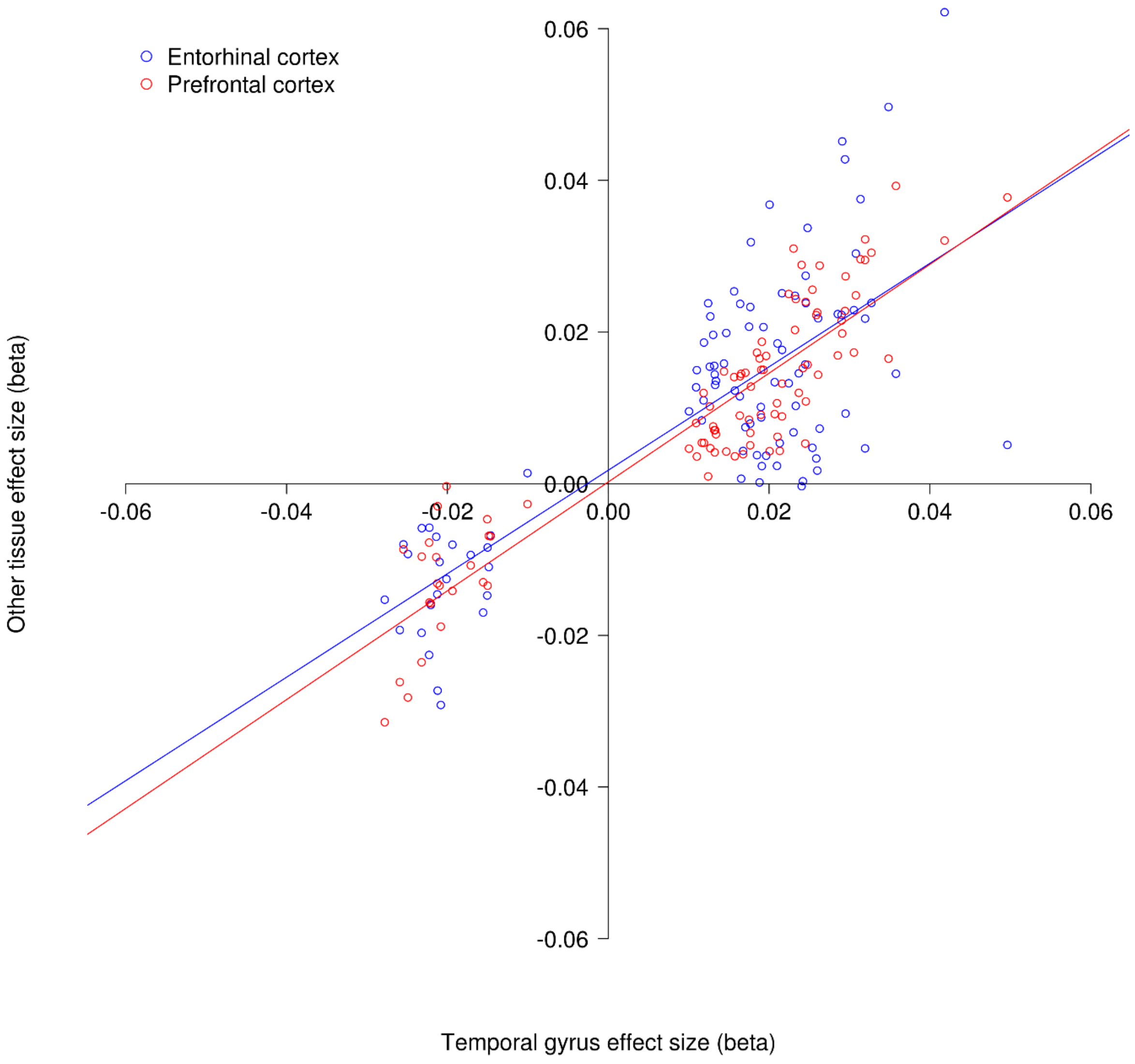

**Supplementary Figure 9: The most significant differentially methylated region (DMR) in the temporal gyrus inverse variance fixed effects meta-analysis (chr8:41469308-41569399) contained 2 probes and resided in the *ANK1* gene (N = 608).** The horizontal red line denotes the Bonferroni significance level of $P < 1.238 \times 10^{-7}$. Red probes represent a positive methylation (beta) effect size (ES) ≥ 0.01, blue probes represent a negative ES ≥ 0.01. Underneath the gene tracks are shown in black with CpG islands in green.

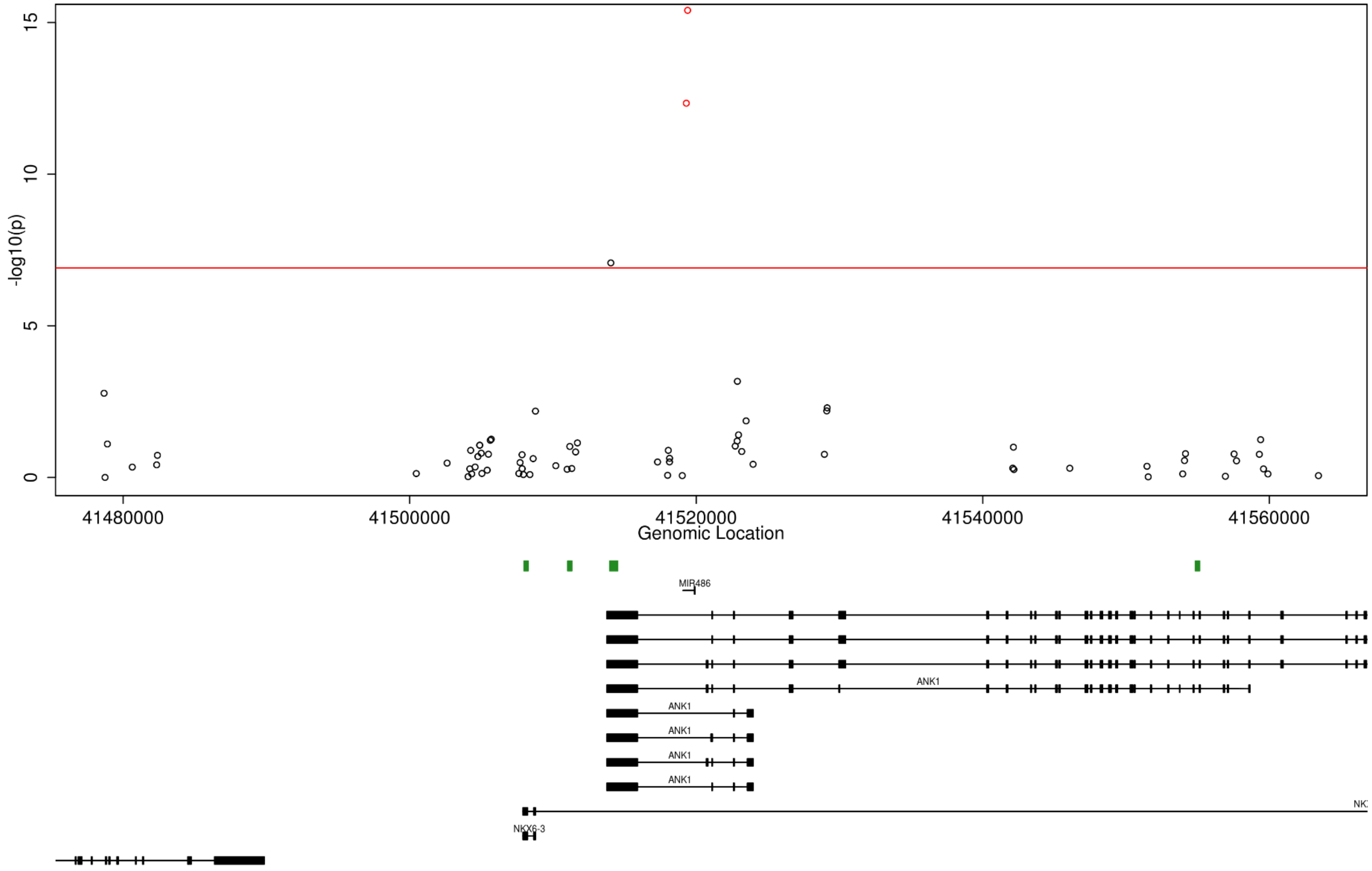

**Supplementary Figure 10: Volcano plot of differentially methylated positions (DMPs) identified in the entorhinal cortex inverse variance fixed effects meta-analysis (N =189).** The X-axis shows methylation (beta) effect size (ES) and the Y-axis shows -log10(p). Gray probes indicate an ES ≥ 0.01, whilst blue probes indicate an ES ≥ 0.01 and a Bonferroni significant p-value ($P < 1.238 \times 10^{-7}$). Exact p-values are provided in Supplementary Data 5.

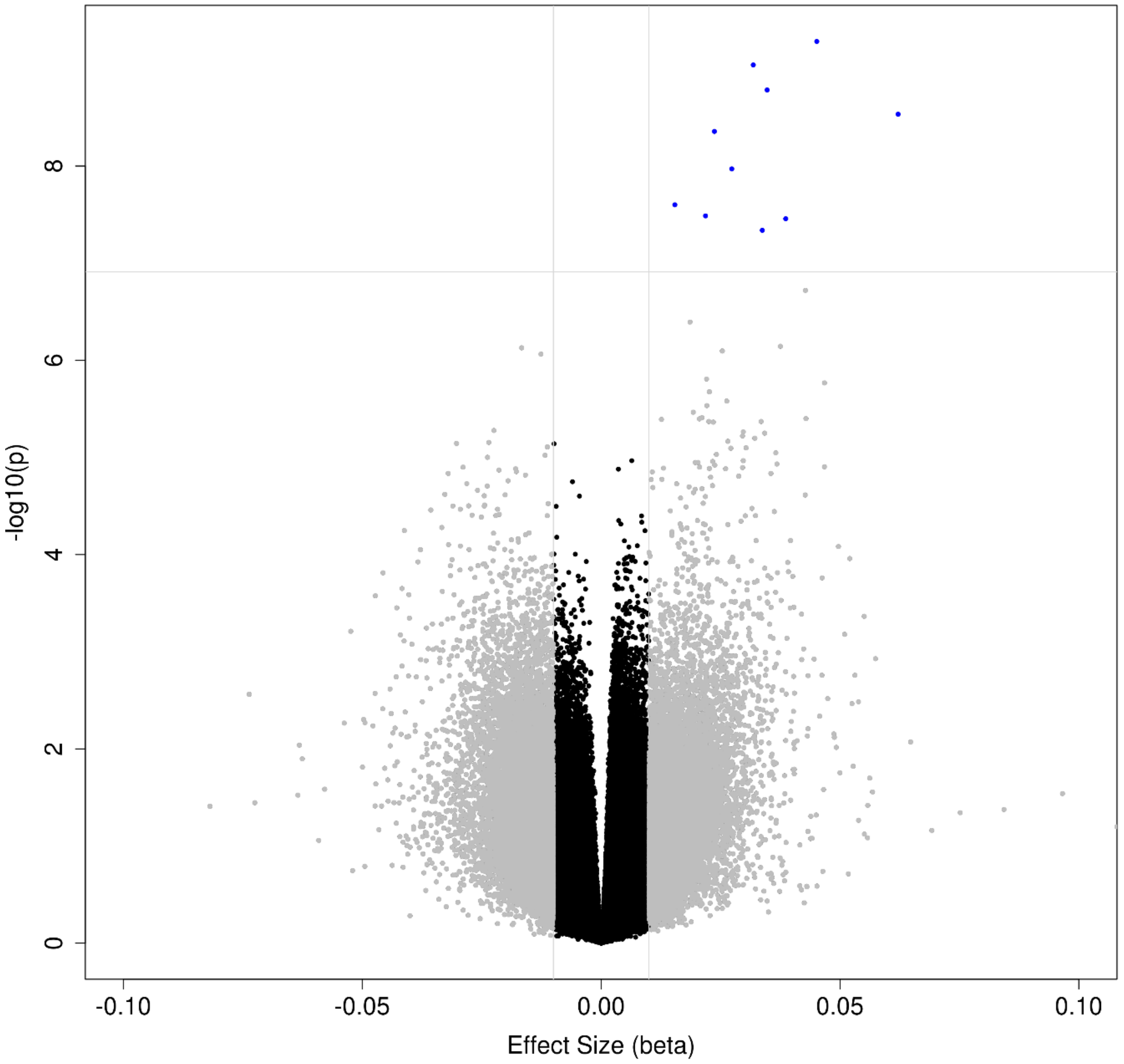

**Supplementary Figure 11: Neuropathology-associated Bonferroni significant differentially methylated positions (DMPs) identified in the entorhinal cortex meta-analysis are replicated in other cortical regions.** The methylation (beta) effect size (ES) of Braak-associated Bonferroni significant DMPs identified in the entorhinal cortex (N = 189) from our inverse variance fixed effect meta-analysis model (X-axis) were significantly correlated with the ES of the same probes in the prefrontal cortex (red; N = 959, r = 0.74, P = 0.01) and temporal gyrus (green; N = 608, r = 0.85, P = 1.82 x $10^{-3}$) (Y-axis).

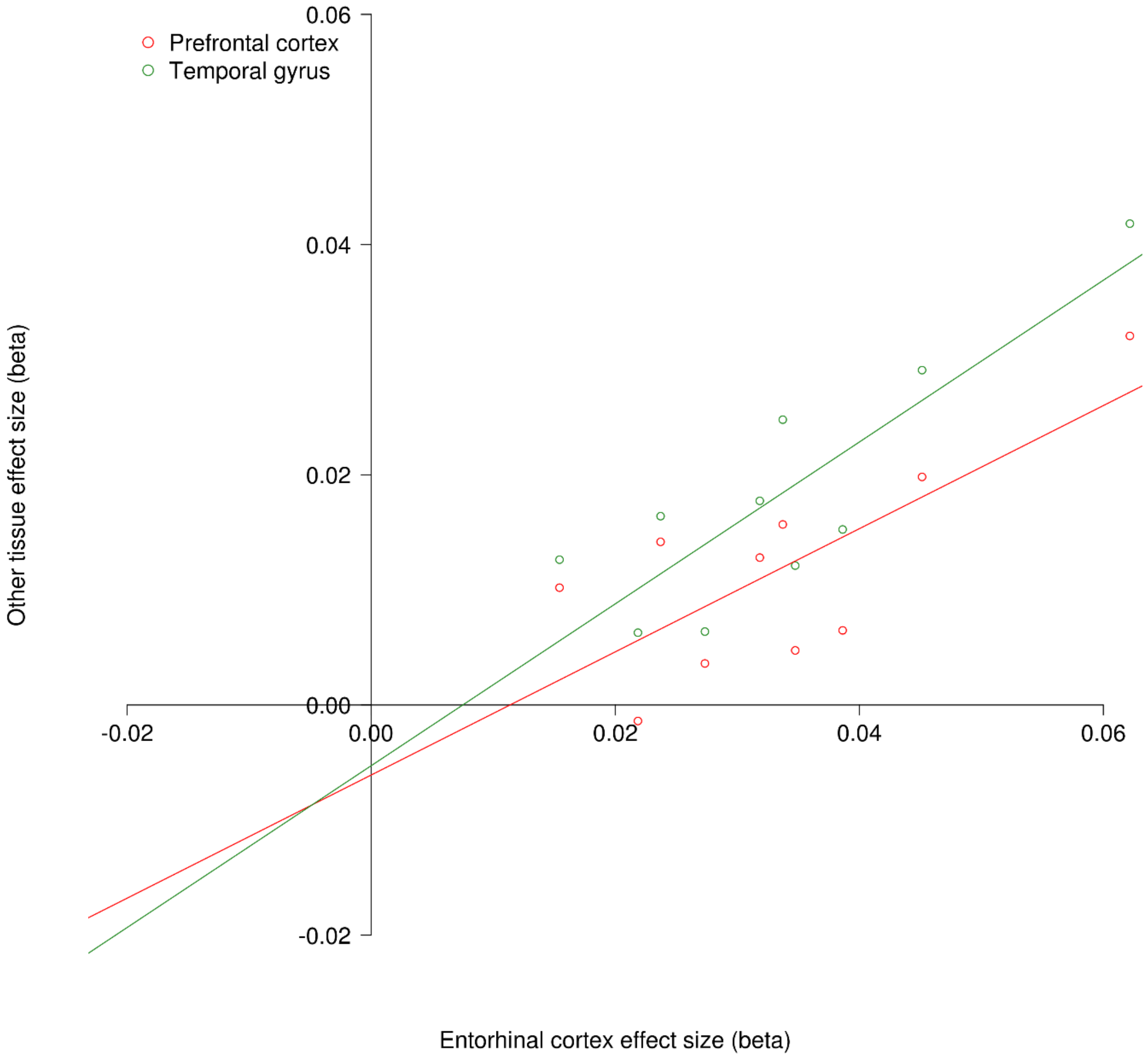

**Supplementary Figure 12: The most significant differentially methylated region (DMR) in the entorhinal cortex inverse variance fixed effects meta-analysis (chr17:74425240-74525402) contained 5 probes and resided in the *RHBDF2* gene (N = 189).** The horizontal red line denotes the Bonferroni significance level of $P < 1.238 \times 10^{-7}$. Red probes represent a positive methylation (beta) effect size (ES) ≥ 0.01, blue probes represent a negative ES ≥ 0.01. Underneath the gene tracks are shown in black with CpG islands in green.

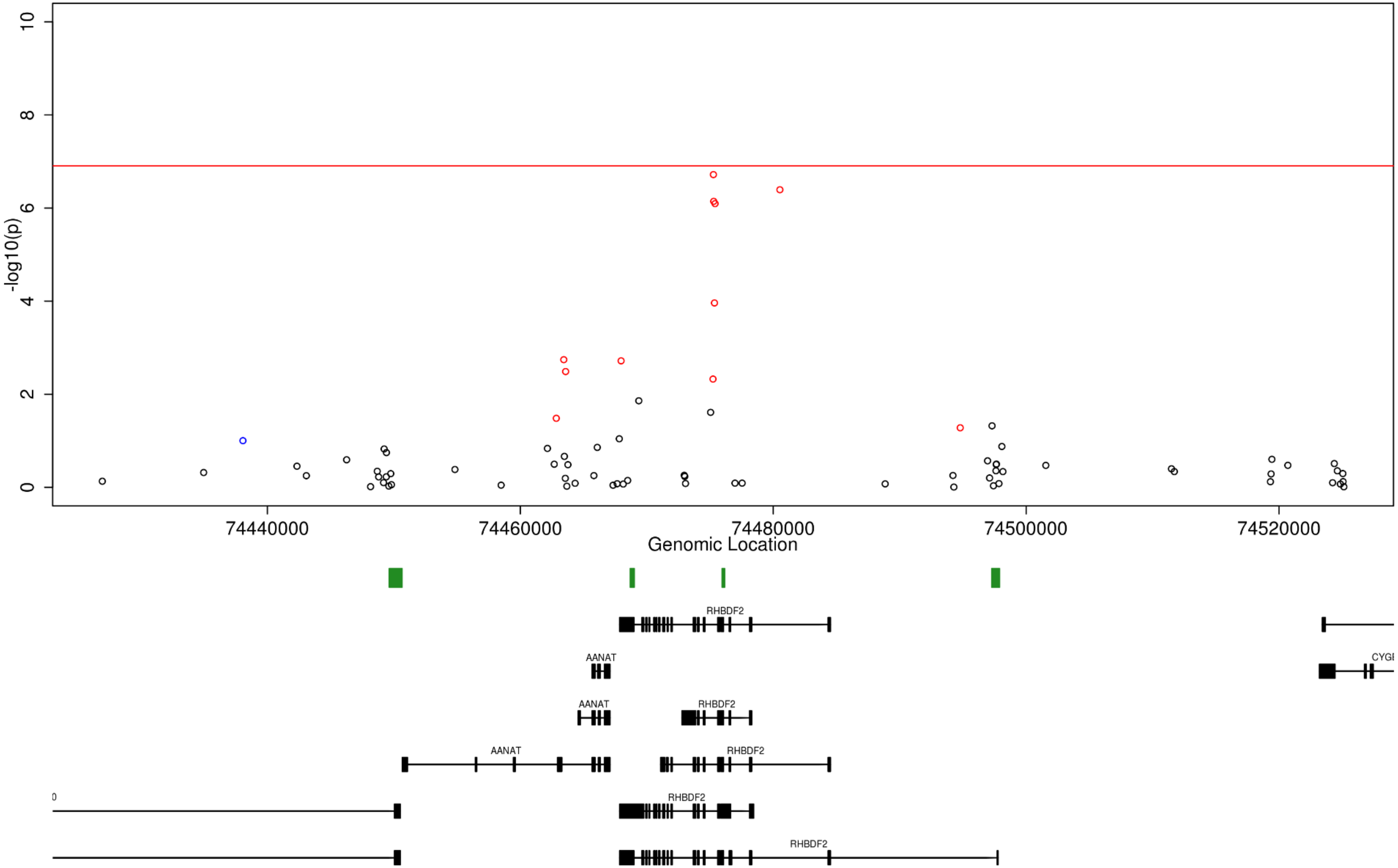

**Supplementary Figure 13: A Manhattan plot for the association of DNA methylation with Braak stage in the cerebellum inverse variance fixed effects meta-analysis (N = 533)**. The X-axis shows chromosome location (Chr1-22) and the Y-axis represents -log10(p-value). The horizontal red line indicates the Bonferroni significance level of P < 1.238 x $10^{-7}$.

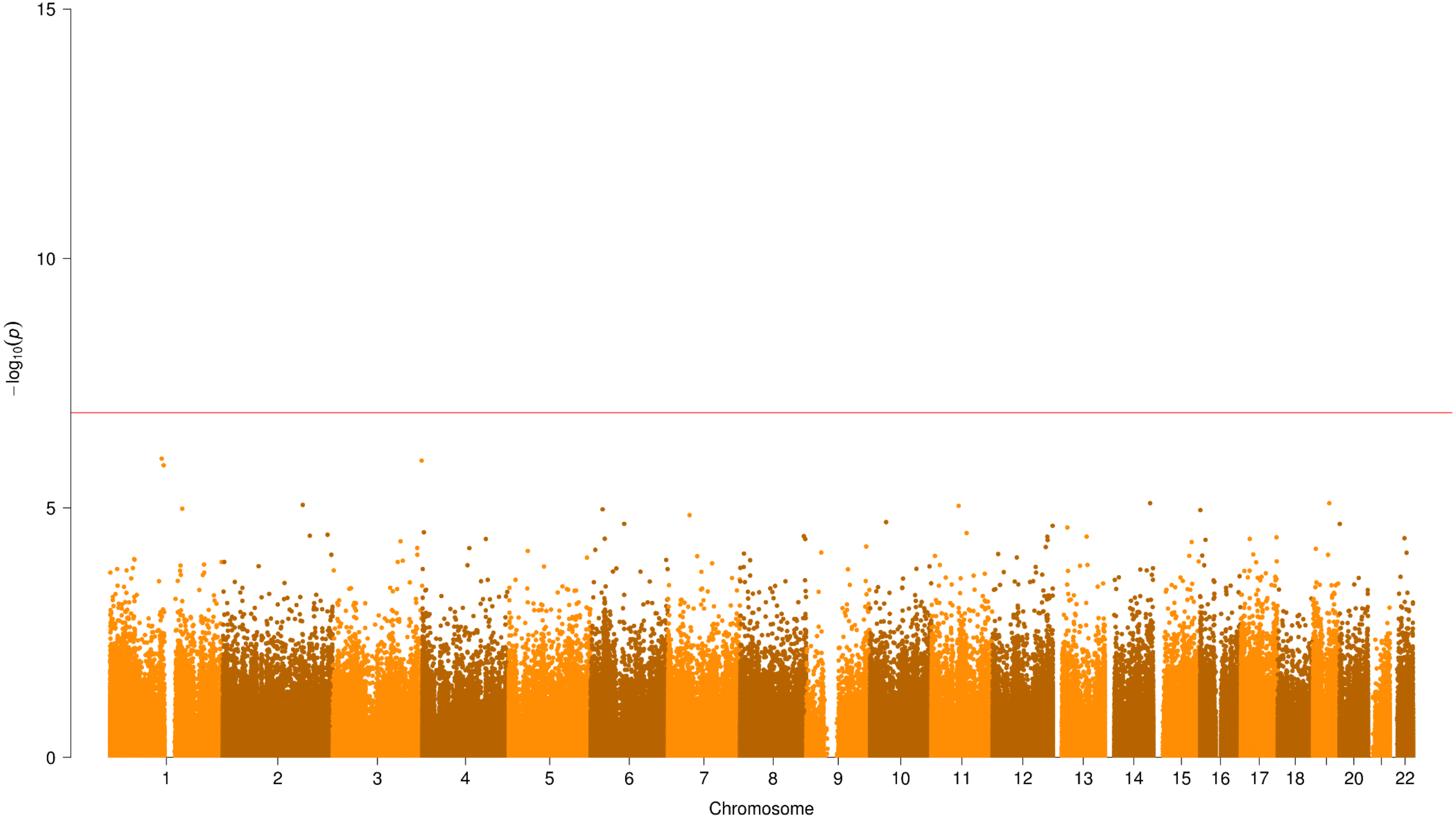

**Supplementary Figure 14: Neuropathology-associated Bonferroni significant differentially methylated positions (DMPs) in the cortex are not seen in the cerebellum.** The methylation (beta) effect size (ES) of the 236 prefrontal cortex (red; N = 959), 95 temporal gyrus (green; N = 608) and ten entorhinal cortex (blue; N = 189) Bonferroni significant Braak-associated DMPs (X-axis) identified in our inverse variance fixed effects meta-analyses were not correlated with the ES of the same probes in the cerebellum (N = 533) (Y-axis).

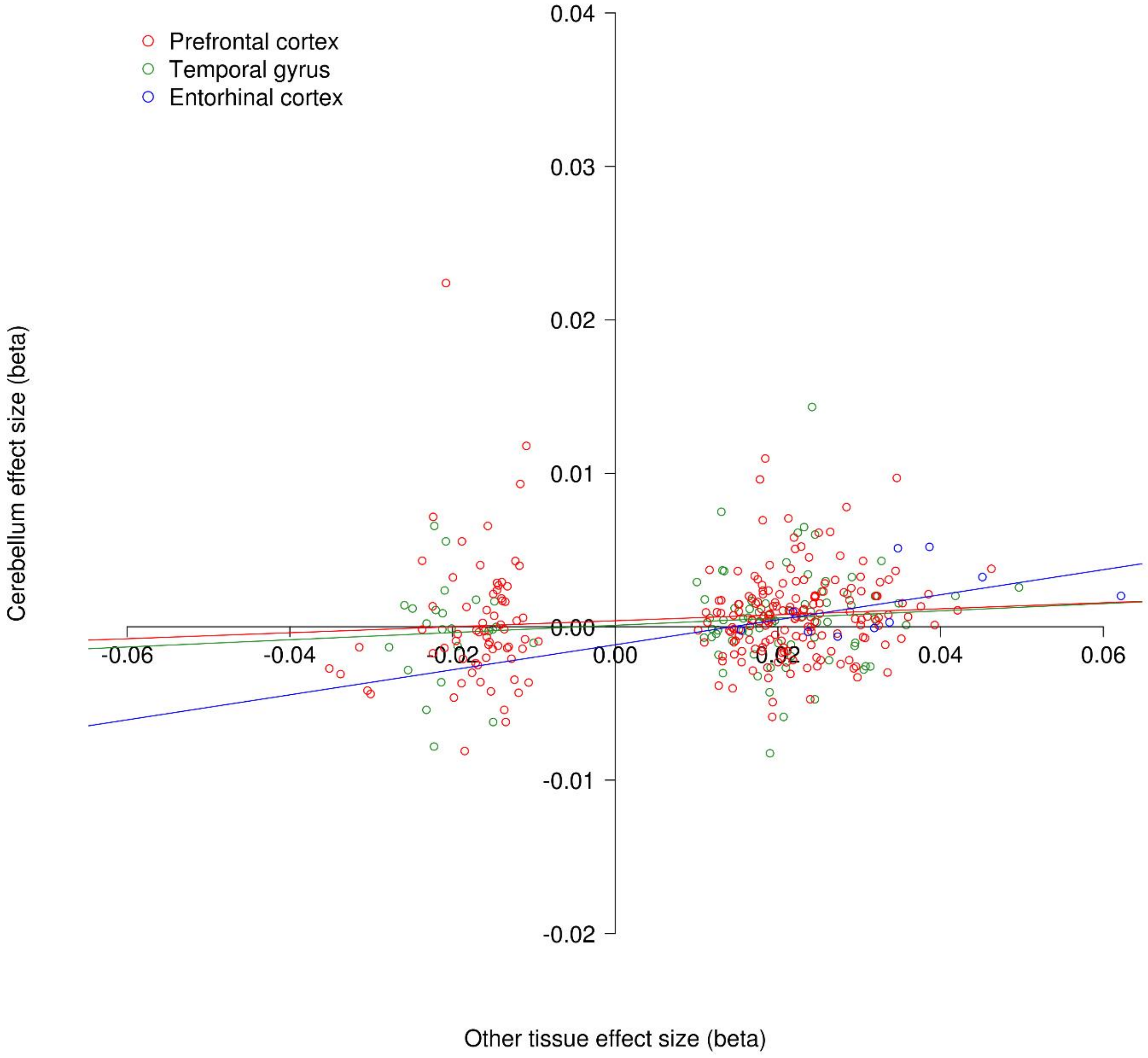

**Supplementary Figure 15: Forest plot of the 18 most significant differentially methylated positions (DMPs) in our cross-cortex inverse variance fixed effects meta-analysis.** The methylation (beta) effect size (ES) is shown in the prefrontal cortex (red; N = 959), temporal gyrus (green; N = 608) and entorhinal cortex (blue; N = 189) for the different cohorts. The X-axis shows the beta ES, with dots representing ES and arms indicating standard error (SE). ES from the intra-tissue meta-analysis using all available individual cohorts are represented by polygons (N = 1,408).

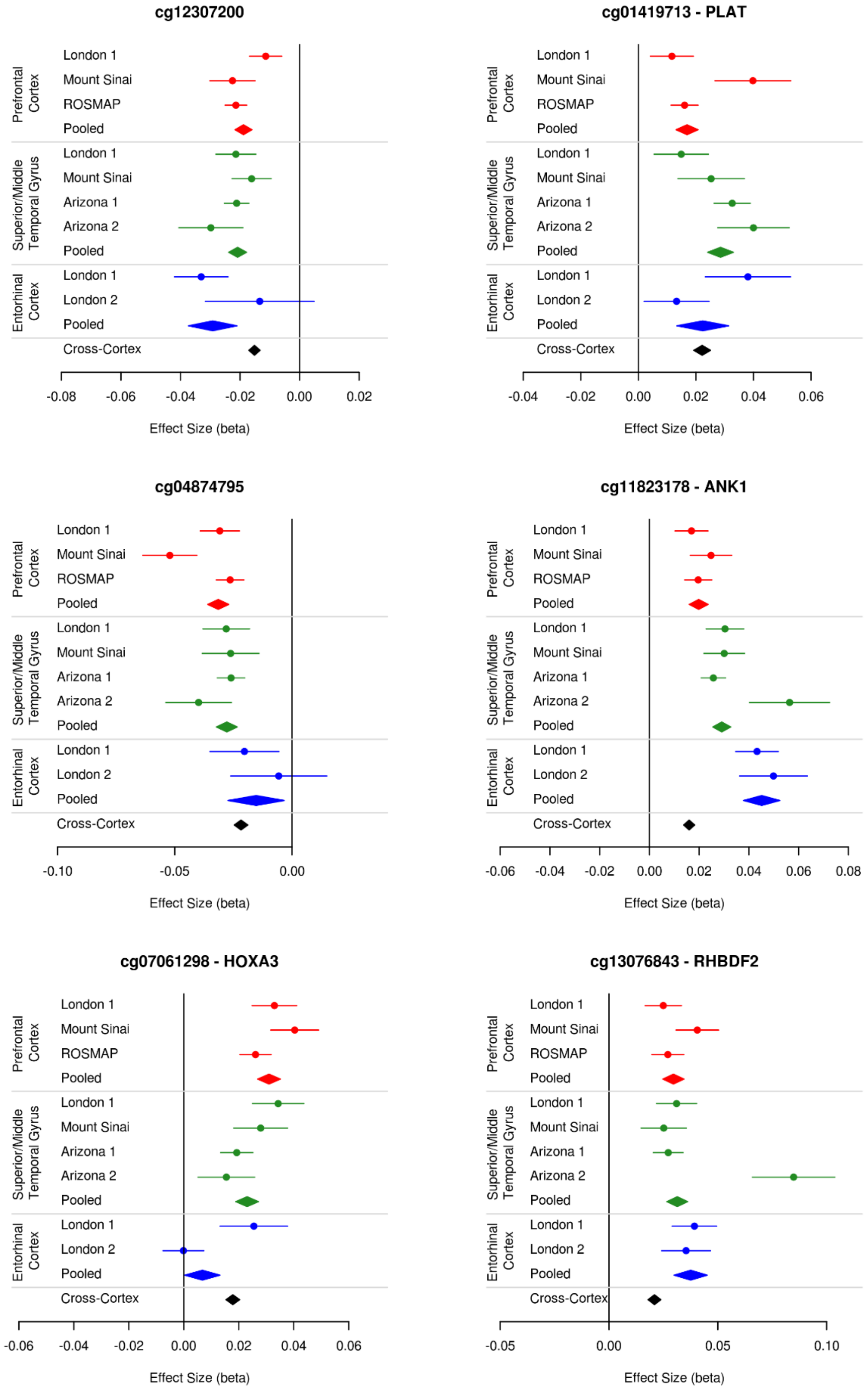

**Supplementary Figure 15 (cont.)**

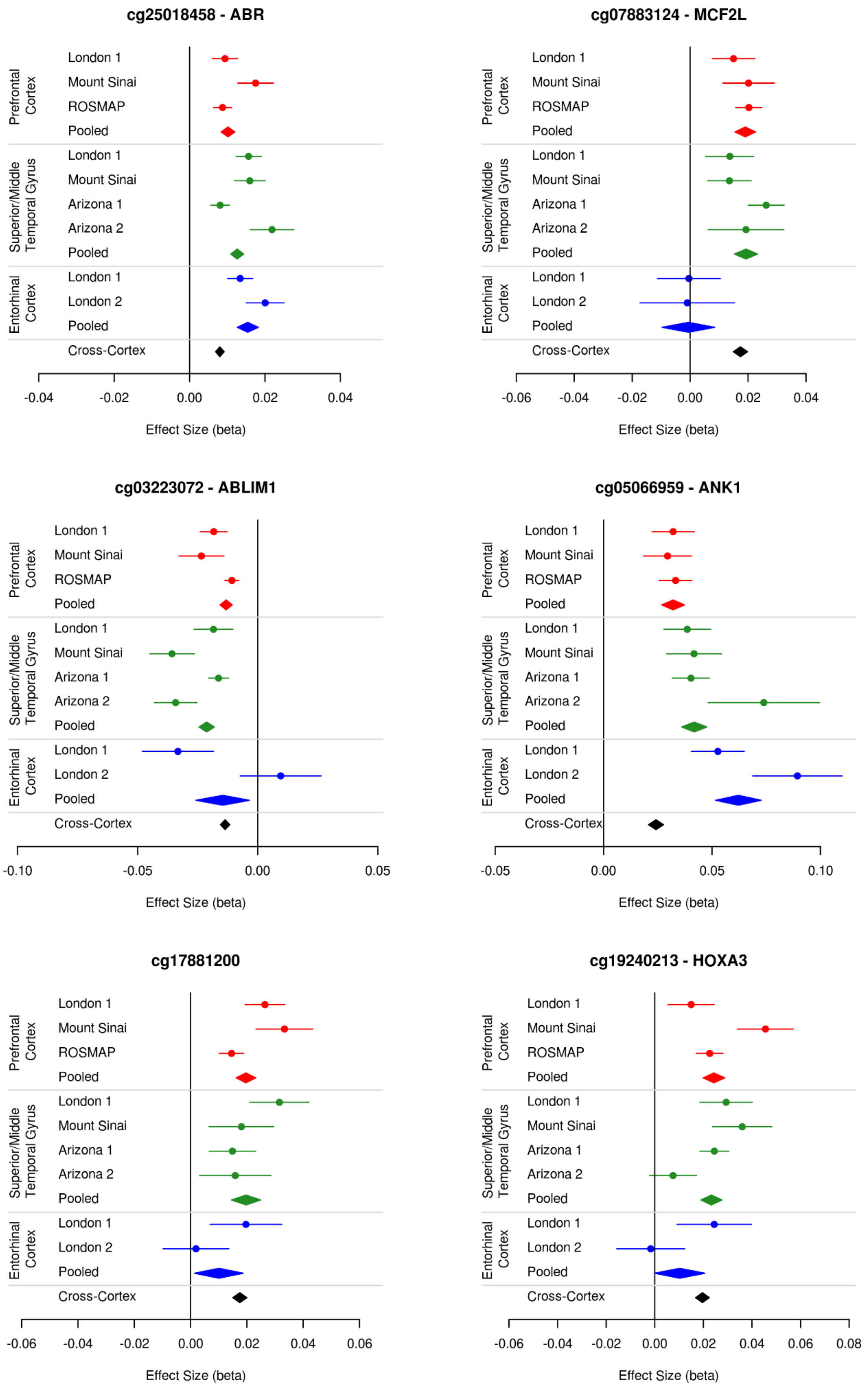

**Supplementary Figure 15 (cont.)**

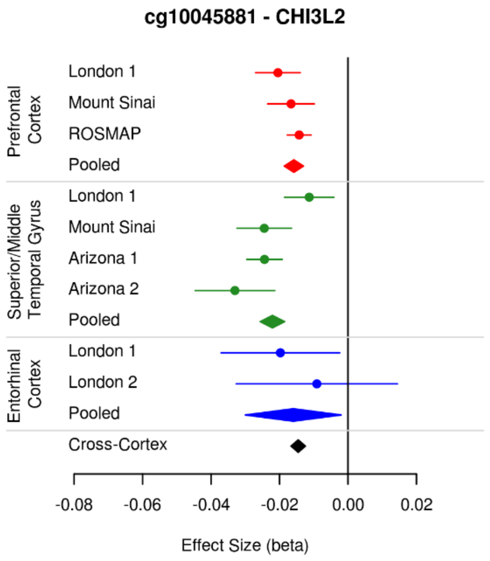


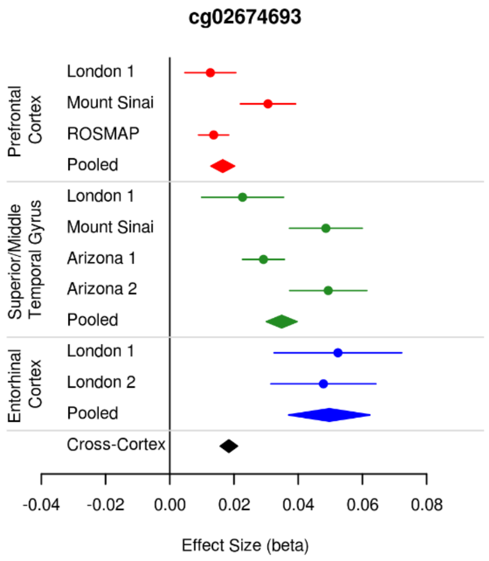


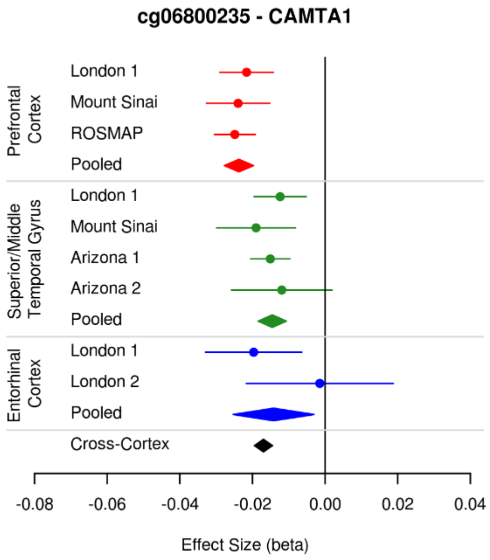


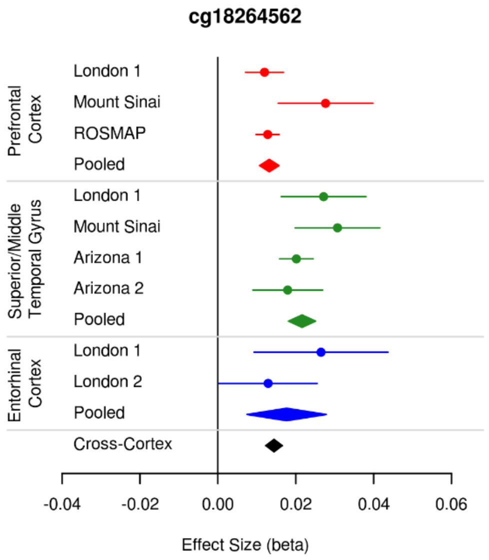


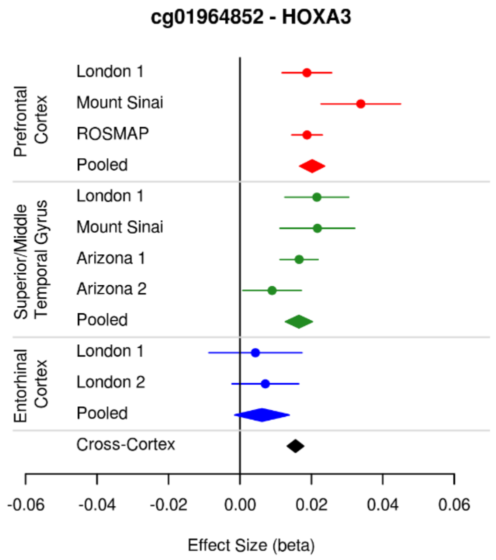


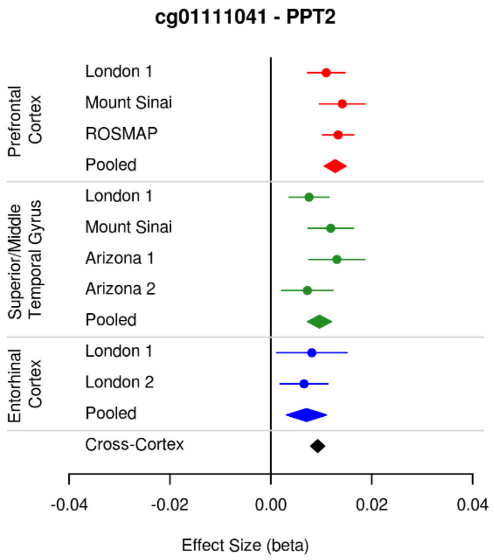

**Supplementary Figure 16: A Venn diagram demonstrating how many of the 220 Bonferroni significant differentially methylated positions (DMPs) in the cross-cortex inverse variance fixed effects meta-analysis were nominally significantly differentially methylated ($P \leq 0.05$) in each of the individual cohorts that contributed to the meta-analysis.**

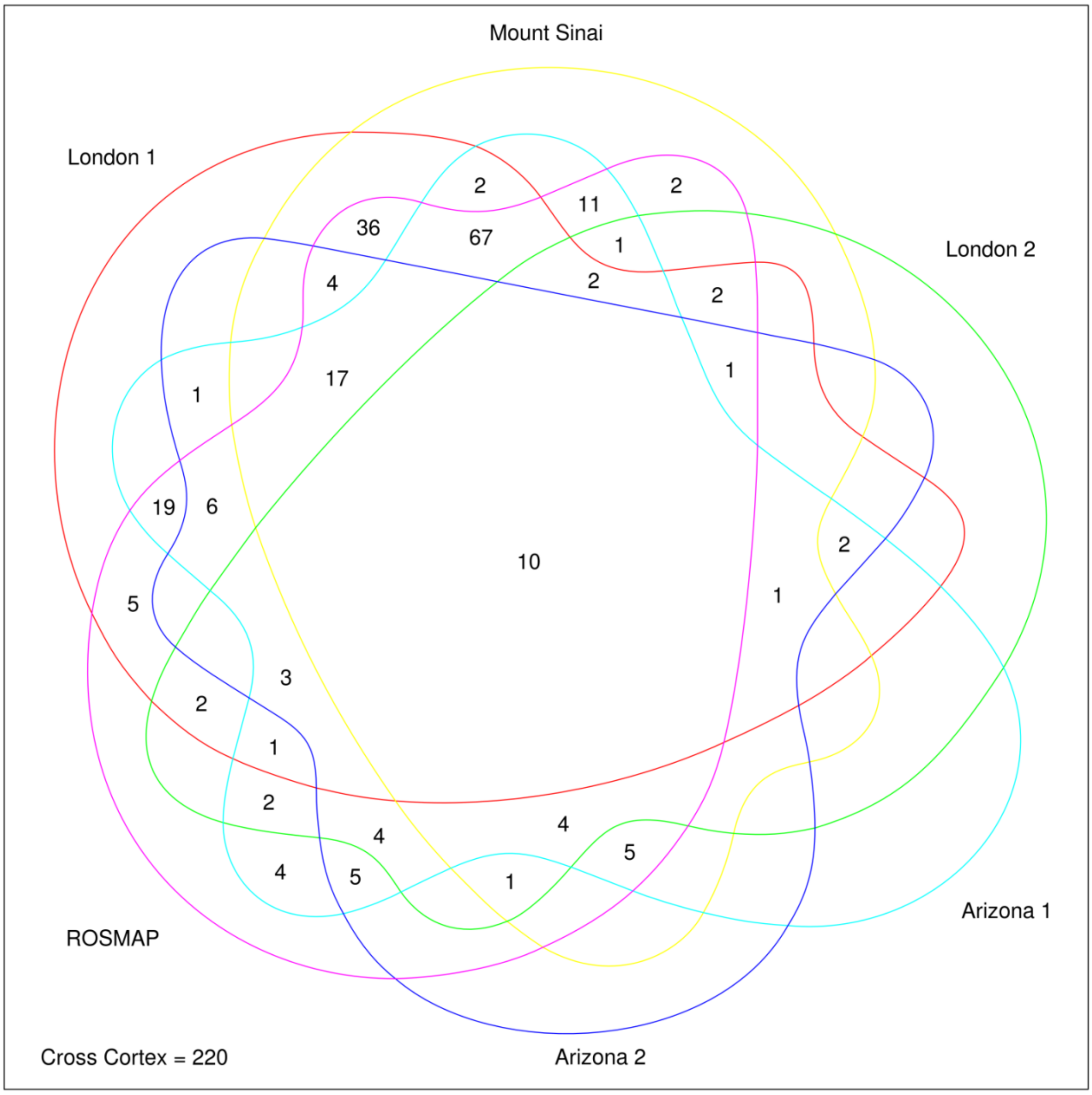

**Supplementary Figure 17: A heat map highlighting the methylation (beta) effect size (ES) and direction of effect for the 220 Bonferroni significant cross-cortex differentially methylated positions (DMPs) across all analyses in all tissues.** Red indicates hypermethylation with higher Braak stage, whilst blue indicates hypomethylation with higher Braak stage

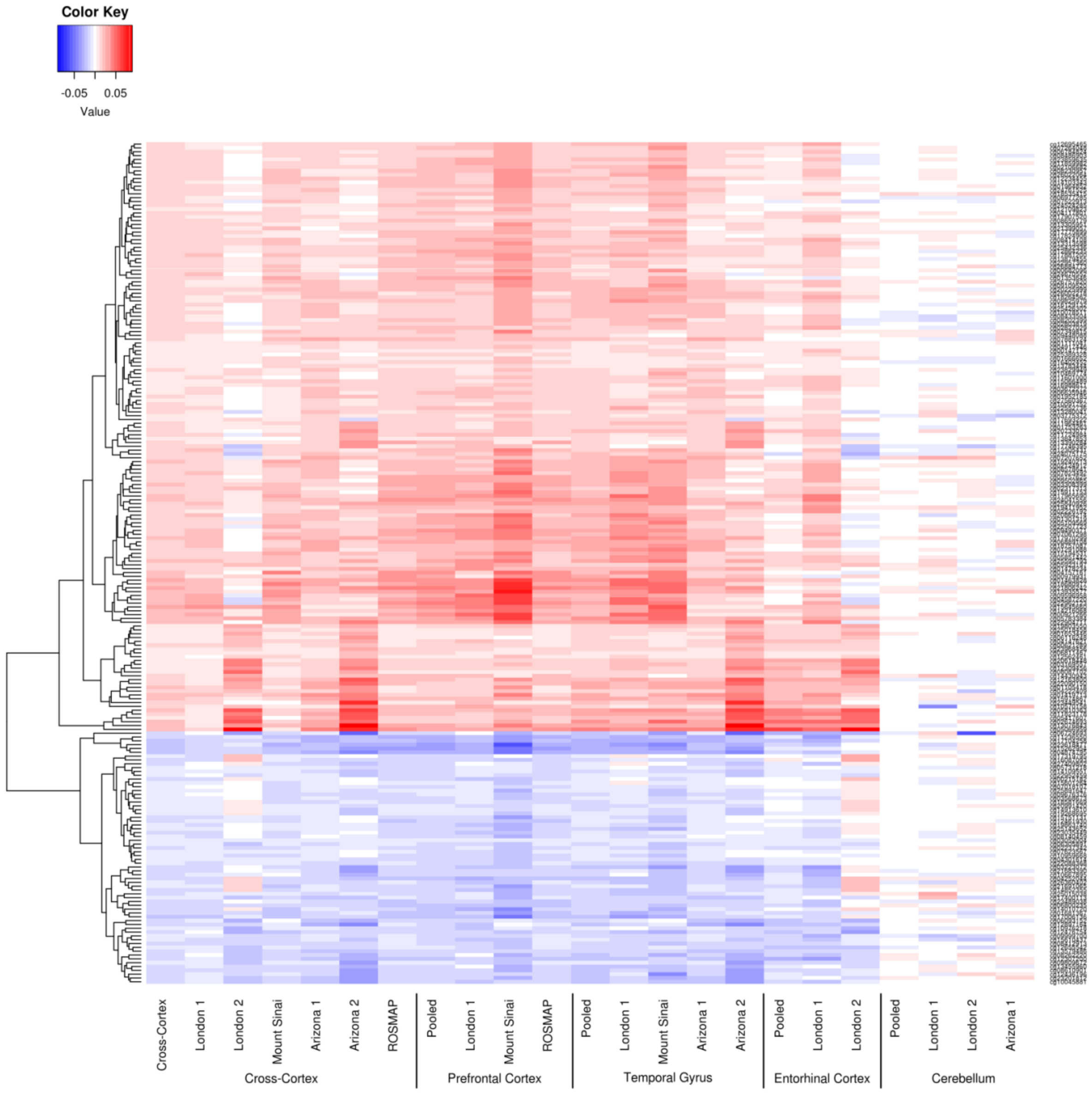

**Supplementary Figure 18: Enrichment of the 220 cross-cortex Bonferroni significant differentially methylated positions (DMPs) in genomic features.** Using a two-sided Fisher's exact test, the cross-cortex Bonferroni significant DMPs were significantly enriched in CpG island shelves and non-CpG islands in the proximal promoter and CpG islands in gene bodies, whilst they were significantly under-represented in CpG islands in the proximal promoter. Key: * = P < 0.05, ** = P < 0.01, *** = P < 0.005, exact p-values can be found in Supplementary Data 11.

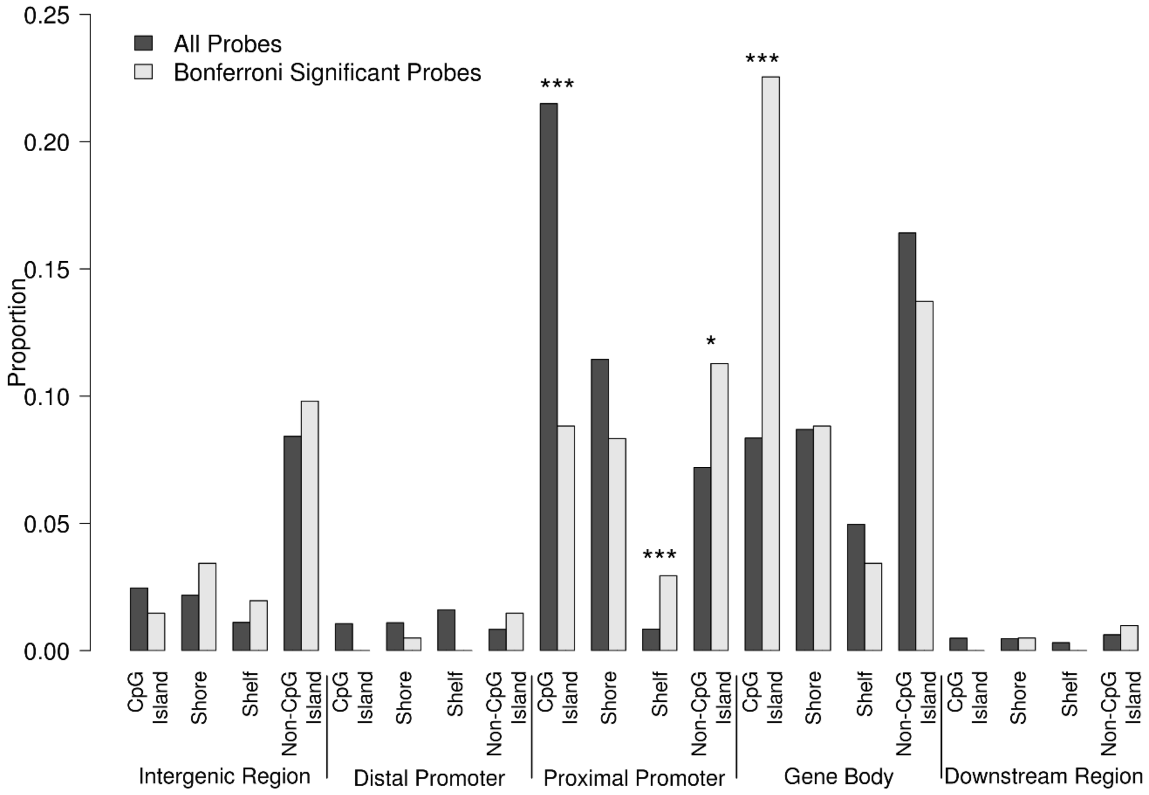

**Supplementary Figure 19: The 220 Bonferroni significant differentially methylated positions (DMPs) from the cross-cortex meta-analysis were significantly under-represented in the 1500bp around the transcription start site, and over-represented in the 1$^{st}$ exon using a two-sided Fisher's exact test.** Shown are the proportions of probes in each transcription related feature for the Bonferroni significant probes and all probes passing quality control. Abbreviations: TSS1500: within 1500bp of transcription start site, TSS200: within 200bp of transcription start site, 5'UTR: 5' untranslated region, 3'UTR: 3' untranslated region. P-values can be found in Supplementary Data 12.

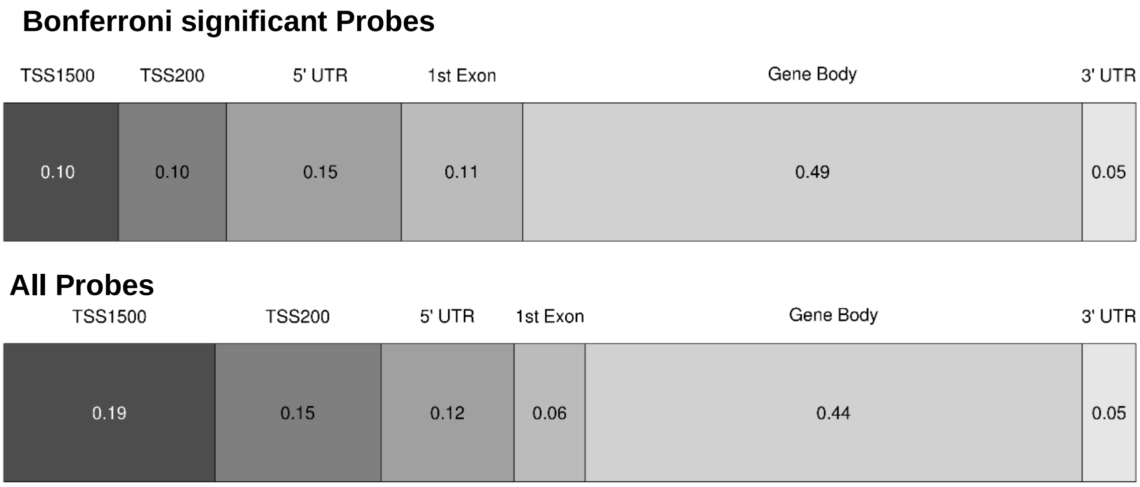

**Supplementary Figure 20: Quantile-Quantile (QQ) plots of the linear mixed-effects analyses.** Shown are QQ plots for (**a**) the prefrontal cortex meta-analysis (N = 959), (**b**) the temporal gyrus meta-analysis (N = 608), (**c**) the entorhinal cortex meta-analysis (N = 189), (d) the cerebellum meta-analysis (N = 533) and (**e**) the cross-cortex meta-analysis (N = 1,408). The X-axis shows expected –log10(p) and Y-axis shows observed –log10(p). A 95% pointwise confidence band (gray area) was computed under the assumption that the p-values were drawn independently from a uniform [0, 1] distribution. All of the analyses had a lambda (λ) < 1.2.

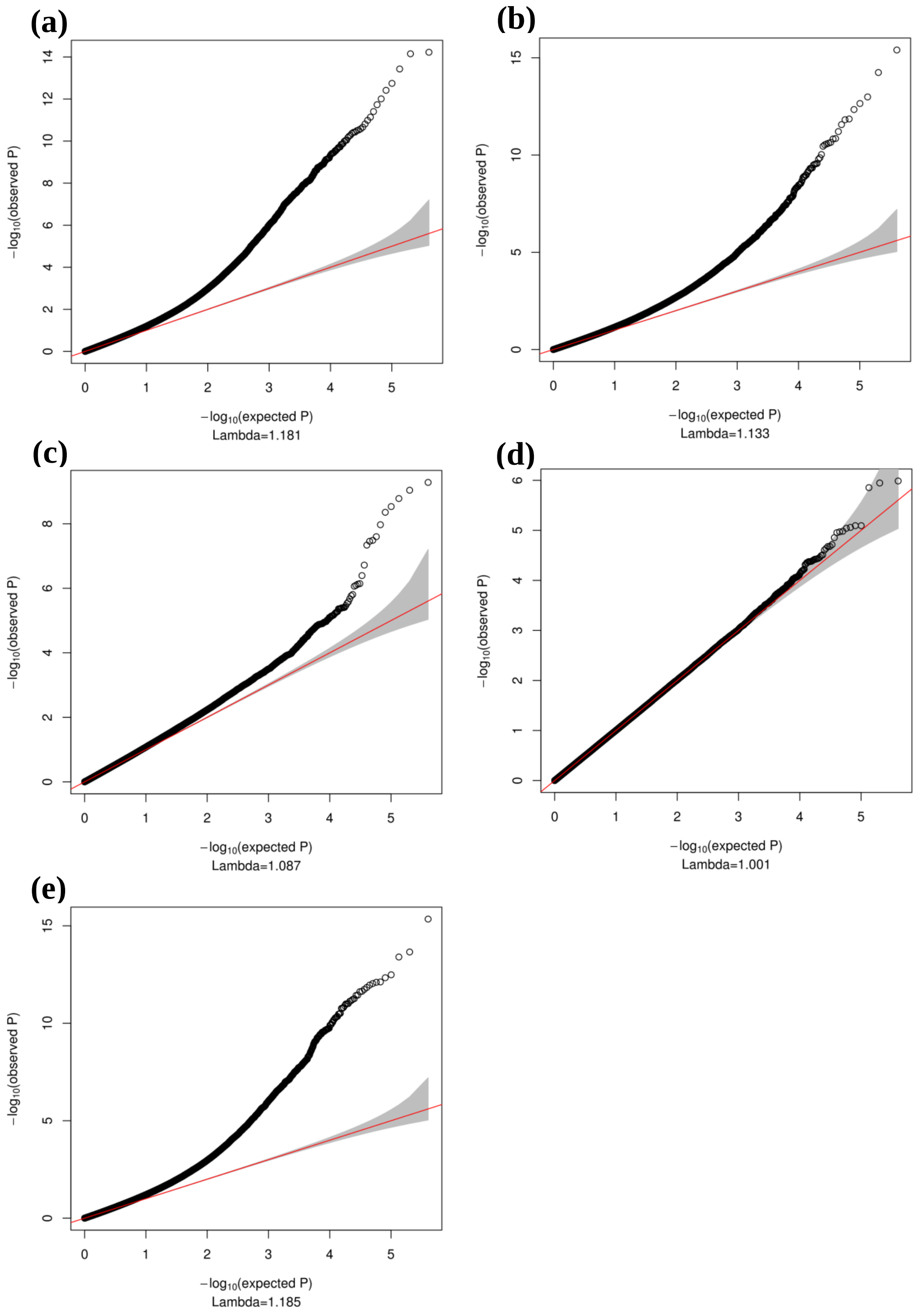
