## Supplementary Data Legends for "A meta-analysis of epigenome-wide association studies in Alzheimer’s disease highlights novel differentially methylated loci across cortex"

**Supplementary Data 1: 236 Bonferroni significant differentially methylated positions (DMPs) were associated with Braak stage in the prefrontal cortex intra-tissue meta-analysis ($P < 1.238 \times 10^{-7}$).** Probe information is provided corresponding to chromosomal location (hg19/GRCh37 genomic annotation), Illumina (UCSC) gene annotation, closest genes with a transcription start site upstream or downstream (from GREAT annotation). Shown for each DMP is the methylation (beta) effect size (ES), standard error (SE) and corresponding unadjusted P value from the prefrontal cortex inverse variance fixed effects meta-analysis and the random effects meta-analysis (N = 959), followed by the heterogeneity statistics (I2 and Heterogeneity P). All ES and SE have been multiplied by six to demonstrate the difference between Braak stage 0 and Braak stage VI samples. Also shown are the ES, SE and unadjusted P value from the inverse variance fixed-effect model in each of the individual cohorts that were used for the prefrontal cortex meta-analysis (London 1 cohort, Mount Sinai cohort, ROS/MAP cohort). In the final column it is highlighted whether that CpG site reached Bonferroni significance individually in the six AD brain EWAS that have been previously published[5-8,10,12,] and which contributed to the current study.

**Supplementary Data 2: Differentially methylated regions (DMRs) significantly associated with Braak stage in the prefrontal cortex intra-tissue meta-analysis.** Shown are all significantly associated regions (Sidak-corrected P value < 0.05) that contain two or more probes from the comb-p regional analysis of the inverse variance fixed-effect intra-tissue meta-analysis of the prefrontal cortex. Shown for DMRs are the chromosomal location (hg19), up/downstream genes from GREAT annotation, number of probes in the significant region and Sidak corrected P value.

**Supplementary Data 3: 95 Bonferroni significant differentially methylated positions (DMPs) were associated with Braak stage in the temporal gyrus intra-tissue meta-analysis ($P < 1.238 \times 10^{-7}$).** Probe information is provided corresponding to chromosomal location (hg19/GRCh37 genomic annotation), Illumina (UCSC) gene annotation, closest genes with a transcription start site upstream or downstream (from GREAT annotation). Shown for each DMP is the methylation (beta) effect size (ES), standard error (SE) and corresponding unadjusted P value from the temporal gyrus inverse variance fixed effects meta-analysis and the random effects meta-analysis (N = 608), followed by the heterogeneity statistics (I2 and Heterogeneity P). All ES and SE have been multiplied by six to demonstrate the difference between Braak stage 0 and Braak stage VI samples. Also shown are the ES, SE and unadjusted P value from the inverse variance fixed-effect model in each of the individual cohorts that were used for the temporal gyrus meta-analysis (London 1 cohort, Mount Sinai cohort, Arizona 1 cohort, Arizona 2 cohort). In the final column it is highlighted whether that CpG site reached Bonferroni significance individually in the six AD brain EWAS that have been previously published[5-8,10,12,] and which contributed to the current study.

**Supplementary Data 4: Differentially methylated regions (DMRs) significantly associated with Braak stage in the temporal gyrus intra-tissue meta-analysis.** Shown are all significantly associated regions (Sidak-corrected P value < 0.05) that contain two or more probes from the comb-p regional analysis of the inverse variance fixed-effect intra-tissue meta-analysis of the temporal gyrus. Shown for DMRs are the chromosomal location (hg19), up/downstream genes from GREAT annotation, number of probes in the significant region and Sidak corrected P value.

**Supplementary Data 5: Ten Bonferroni significant differentially methylated positions (DMPs) were associated with Braak stage in the entorhinal cortex intra-tissue meta-analysis ($P < 1.238 \times 10^{-7}$).** Probe information is provided corresponding to chromosomal location (hg19/GRCh37 genomic annotation), Illumina (UCSC) gene annotation, closest genes with a transcription start site upstream or downstream (from GREAT annotation). Shown for each DMP is the methylation (beta) effect size (ES), standard error (SE) and corresponding unadjusted P value from the entorhinal cortex inverse variance fixed effects meta-analysis and the random effects meta-analysis (N = 189), followed by the heterogeneity statistics (I2 and Heterogeneity P). All ES and SE have been multiplied by six to demonstrate the difference between Braak stage 0 and Braak stage VI samples. Also shown are the ES, SE and unadjusted P value from the inverse variance fixed-effect model in each of the individual cohorts that were used for the entorhinal cortex meta-analysis (London 1 cohort, London 2 cohort). In the final column it is highlighted whether that CpG site reached Bonferroni significance individually in the six AD brain EWAS that have been previously published[5-8,10,12,] and which contributed to the current study.

**Supplementary Data 6: Differentially methylated regions (DMRs) significantly associated with Braak stage in the entorhinal cortex intra-tissue meta-analysis.** Shown are all significantly associated regions (Sidak-corrected P value < 0.05) that contain two or more probes from the comb-p regional analysis of the inverse variance fixed-effect intra-tissue meta-analysis of the entorhinal cortex. Shown for DMRs are the chromosomal location (hg19), up/downstream genes from GREAT annotation, number of probes in the significant region and Sidak corrected P value.

**Supplementary Data 7: 220 Bonferroni significant differentially methylated positions (DMPs) were associated with Braak stage in the cross-cortex meta-analysis ($P < 1.238 \times 10^{-7}$).** Probe information is provided corresponding to chromosomal location (hg19/GRCh37 genomic annotation), Illumina (UCSC) gene annotation, closest genes with a transcription start site upstream or downstream (from GREAT annotation). Shown for each DMP is the methylation (beta) effect size (ES), standard error (SE) and corresponding unadjusted P value from the inverse variance fixed effects meta-analysis and the random effects meta-analysis in the cross-cortex data (N = 1,408), followed by the heterogeneity statistics (I2 and Heterogeneity P). All ES and SE have been multiplied by six to demonstrate the difference between Braak stage 0 and Braak stage VI samples. Also shown are the ES, SE and unadjusted P value from the inverse variance fixed-effect model in each of the individual cohorts that were used for the meta-analysis (London 1 cohort, London 2 cohort, Mount Sinai cohort, Arizona 1 cohort, Arizona 2 cohort, ROS/MAP cohort). In the penultimate column it is highlighted whether that CpG site reached Bonferroni significance individually in the six AD brain EWAS that have been previously published[5-8,10,12,] and which contributed to the current study. In the final column it is highlighted whether that CpG reached nominal significance (P <0.05) in all six discovery cohorts used for the meta-analysis.

**Supplementary Data 8: Gene Ontology pathway analysis on the 121 annotated genes corresponding to the 220 Bonferroni significant differentially methylated positions (DMPs) from the inverse variance fixed effects cross-cortex meta-analysis.** Shown is the gene ontology ID, a description of the pathway, the type of pathway, the number of genes annotated to the pathway, the number of genes annotated to the Bonferroni significant DMPs in that pathway, the corresponding unadjusted P value, the false discovery rate (FDR) adjusted P value (Q value) and the genes in the test list that are present in that pathway. Pathways are only shown in the table if unadjusted P ≤ 0.05.

**Supplementary Data 9: Identification of 165 independent signals in the 220 Bonferroni significant differentially methylated positions (DMPs) that were identified in the cross-cortex meta-analysis.** We collapsed the 220 Bonferroni significant loci into 165 independent, non-highly correlated signals using a threshold of r < 0.6 over 1mb using a method for identifying SNPs in linkage disequilibrium (LD)[14] and implementing this in the ROSMAP data (as the largest single tissue dataset). Shown are the collapsed regions where ≥ 2 probes were present originally and have been reduced to fewer independent signals.

**Supplementary Data 10: Differentially methylated regions (DMRs) significantly associated with Braak stage in the cross-cortex meta-analysis.** Shown are all significantly associated regions (Sidak-corrected P value < 0.05) that contain two or more probes from the comb-p regional analysis of the inverse variance fixed-effect cross-cortex meta-analysis. Shown for DMRs are the chromosomal location (hg19), up/downstream genes from GREAT annotation, number of probes in the significant region and Sidak corrected P value.

**Supplementary Data 11: Enrichment analysis of genomic features annotated to the 220 Bonferroni significant differentially methylated positions (DMPs) in the cross-cortex meta-analysis.** A two-sided Fisher's exact test was used to determine whether Bonferroni significant DMPs were enriched in particular genomic features based on Slieker annotation. Shown are genomic region and feature, total number of probes passing QC corresponding to each feature, number of Bonferroni significant probes corresponding to each feature, odds ratio (OR) and the corresponding P value. Shown in bold are significant enrichments.

**Supplementary Data 12: Enrichment analysis of transcription related features annotated to the 220 Bonferroni significant differentially methylated positions (DMPs) in the cross-cortex meta-analysis.** A two-sided Fisher's exact test was used to determine whether Bonferroni significant DMPs were enriched in particular transcription related features based on Illumina annotation. Shown are feature, total number of probes passing QC corresponding to each feature, number of Bonferroni significant probes corresponding to each feature, and the corresponding odds ratio (OR) and P value. Shown in bold are significant enrichments. Abbreviations: TSS1500: within 1500bp of transcription start site, TSS200: within 200bp of transcription start site, 5'UTR: 5' untranslated region, 3'UTR: 3' untranslated region.

**Supplementary Data 13: The subset of 110 probes from the 220 Bonferroni significant cross-cortex differentially methylated positions (DMPs) that can predict whether a sample has high or low levels of pathology based on DNA methylation levels.** Shown are the probe ID, genomic location, Illumina gene annotation and coefficient. Data is ordered by the largest contribution to the classifier (i.e. absolute coefficient value).

**Supplementary Data 14: Variance in Braak pathology can be explained by DNA methylation signatures.** An elastic net penalized regression analysis model was used in the discovery (training) dataset to identify 110 probes (see Supplementary Data 13) that can explain variance in pathology between samples with low pathology (Braak 0-II: control) compared to high pathology (Braak V-VI: AD). Training data consisted of all Braak 0-II and Braak V-VI samples (Braak 0-II: N = 407, Braak V-VI: N = 589). Subsequently this was then tested for performance in 38 samples from the Munich replication prefrontal cortex cohort (Braak 0-II: N = 9, Braak V-VI: N = 29) and 454 samples from the BDR replication prefrontal cortex cohort (Braak 0-II: N = 196, Braak V-VI: N 258). Abbreviations: area under curve (AUC), confidence interval (CI).

**Supplementary Data 15: Eighteen of the Bonferroni significant cross-cortex differentially methylated positions (DMPs) represented methylation quantitative trait loci (mQTLs).** Using the xQTL resource fron Ng et al.[17] we searched for the 220 Bonferroni significant cross-cortex DMPs in the list of CpGs significantly associated with single nucleotide polymorphisms (SNPs) from the xQTL resource. We identified 200 mQTLs relating to 18 of the Bonferroni significant DMPs that represent significant mQTLs. Shown are the Illumina (UCSC) Gene Annotation for the 450K Probe, the SNP within that genomic region that represents a cis-mQTL, the genomic location of the SNP, the Spearman Rho and P value for the mQTL.

**Supplementary Data 16: Analysis of probes located in known Alzheimer's disease (AD) linkage disequilibrium (LD) blocks nominated from a recent genome-wide association study (GWAS) meta-analysis.** Of the 24 blocks reported by Kunkle et al.[18] of these contained > 1 CpG site on the 450K array. We used Brown's method to combine P values for CpGs in these AD risk LD blocks. Reported are the best individual DMP value within the region, the combined P value across all the CpGs and the Bonferroni-adjusted combined P value for the cross-cortex, prefrontal cortex, temporal gyrus and entorhinal cortex meta-analyses.

**Supplementary Data 17: Table of surrogate variables used for modelling data.** Shown are the number of surrogate variables detected in each individual analysis and then the number of these added to each model, and the corresponding lambda. * The methylation data on the cerebellum samples for Arizona 1 and Arizona 2 were generated in the same experiment, so these cerebellum samples were analyzed as a single dataset.
